# Challenges in detecting ecological interactions using sedimentary ancient DNA data

**DOI:** 10.1101/2024.08.16.608343

**Authors:** Fiona Margaret Callahan, Jacky Kaiyuan Li, Rasmus Nielsen

## Abstract

With increasing availability of ancient and modern environmental DNA technology, whole-community species occurrence and abundance data over time and space is becoming more available. Sedimentary ancient DNA data can be used to infer associations between species, which can generate hypotheses about biotic interactions, a key part of ecosystem function and biodiversity science. Here, we have developed a realistic simulation to evaluate five common methods from different fields for this type of inference. We find that across all methods tested, false discovery rates of inter-species associations are high under simulation conditions where the assumptions of the methods are violated in a variety of ecologically realistic ways. Additionally, we find that for more realistic simulation scenarios, with sample sizes that are currently realistic for this type of data models are typically unable to detect interactions better than random assignment of associations. Different methods perform differentially well depending on the number of taxa in the dataset. Some methods (SPIEC-EASI, SparCC) assume that there are large numbers of taxa in the dataset, and we find that SPIEC-EASI is highly sensitive to this assumption while SparCC is not. Additionally, we find that for many methods, default calibration can result in high false discovery rates. We find that for small numbers of species, no method consistently outperforms logistic and linear regression, indicating a need for further testing and methods development.

## Introduction

Having a better understanding of how and why ecological communities change over time facilitates informed decisions regarding management and environmental protection [1–3]. One of the key components of this is to understand the influence of species on each other as communities assemble and during periods of environmental change [4, 5]. Inclusion of species interactions in ecological modeling changes predictions about the ecological effects of climate change [4, 6], extinction events [5], and the dynamics of whole ecosystems [2]. Species distribution data up to this point has been largely limited to spatial data, or data over short time periods relative to many of the ecological processes at play [1]. However, as sedimentary ancient DNA (sedaDNA) data becomes more available, it is becoming possible to survey populations across large spatial and temporal extents, and to simultaneously capture data across a wide range of taxa [1, 3]. This data presents the opportunity to study how different taxa have co-occurred over large spatiotemporal scales, and to make inferences about association networks among species [1, 2].

Several categories of methods have been used to infer associations between species using data from various proxies for species presence or abundance, including sedaDNA [7–9]. These associations may arise from interactions among the species or from other sources such as shared responses to the environment [9, 10]. One of the most popular correlative models for spatiotemporal inference using presence-absence data are referred to as species distribution models (SDM) or joint species distribution models (JDSM) [11]. These are generalized linear mixed models with either a logit or probit link function and can include a random effect that accounts for spatiotemporal autocorrelation [12–14]. This class of models is highly varied and can include components that account for a variety of factors, but in most cases, they are designed for very few species in a single study and therefore may not scale to the number of species often seen in sedaDNA data sets (except see [15]). An important feature of (some of) these methods is that they can account for autocorrelation between samples in space and time [12]. Failing to account for this can result in high rates of false inference [16]. However, there are still some potentially important dynamics of this data type not accounted for by these models. Many JSDMs, including those examined here, do not account for uncertainty in covariates and time points, non-linear effects, and false detections of species (but see [17, 18]).

Though SDMs have primarily been used to detect correlations between species and their abiotic environment, some have also been used to detect both biotic and abiotic interactions [12]. Wang, et al. used a SDM method in a study that investigated ecological interactions over the last fifty thousand years in the Arctic using sedaDNA data [12]. They make inferences about the ecological dynamics between humans and megafauna, plants and megafauna, and co-occurrence of different species over a large spatiotemporal scale [12]. Here, we will refer to the methods they used as SDM-INLA. Another study implemented a JSDM for spatiotemporal ordinal abundance data using a Markov Chain Monte Carlo (MCMC) algorithm [14]. They modeled the co-occurrence patterns of several frog species and of several Eucalypts, concluding that frog species had positive residual correlation not accounted for by measured environmental variables, whereas the Eucalypts had negative residual co-occurrence patterns after accounting for the measured environmental variables [14]. We will refer to this method here as JSDM-MCMC.

*Network analysis* methods are another set of correlative methods for detecting associations between taxa using sedaDNA data [19]. They are most often applied to microbial data, though not exclusively [20]. These methods include many different approaches, but in general they use aspects of mathematical network theory in inference and interpretation of associations. These methods are often used for time series data (i.e. a single sediment core subsampled vertically, representing sampling through time), although the methods considered here do not explicitly model time or space [8, 9, 21]. SPIEC-EASI and SparCC [8, 21] use sedaDNA read abundances per taxon as input. EcoCopula [9], which is not explicitly designed for sedaDNA, can accommodate count, biomass, or presence-absence data, and incorporates covariates and species interactions. These *network analysis* methods have been used to investigate microbial associations in the human gut microbiome [8], link microbial network complexity to ecosystem functionality [22], and investigate associations among a broad range of taxa in a marine ecosystem over a period of vast environmental change [20]. In the microbial context, DNA sequences are generally not assigned to species or taxa. Instead, reads are grouped into operational taxonomic units (OTUs) or amplicon sequence variants (ASVs) using sequence similarity [23, 24]. On the other hand, in studies involving plants and megafauna, sequences are often assigned at a species or genus level [12]. While we recognize that there are differences between these data types in real data, since all data are simulated in this study, we will use species, taxon, OTU, and ASV interchangeably here.

The associations inferred by these methods may arise from 1) direct interactions between species such as trophic, mutualistic, or competitive interactions, 2) indirect interactions such as similar responses to the same environment, or 3) falsely inferred associations. If species are associated through an unmeasured aspect of the environment or through an unmeasured species, associations inferred through correlative models may not be causal [10]. Therefore, we do not expect all inferred associations to be causal, but rather consider the aim of these models to be to generate hypotheses about causal interactions, which can then later be tested experimentally [10]. Additionally, false inferences will always be expected to occur at some low rate due to stochastic effects, but violations of modeling assumptions such as non-equilibrium dynamics, differing statistical distributions of the data and the models, or uncertainty in covariates can cause much higher rates of false inference [10, 25, 26].

One of the difficulties of inferring associations between species in this setting is that the number of possible pairs of taxa scales with the square of the number of taxa in the dataset [27]. In studies focusing on megafauna or plants the number of taxa is most often in the tens [12, 14], but in microbiome studies, the number of taxa can be in the hundreds [8, 22]. This means that the number of parameters that are being inferred (interactions between pairs of species) can be much larger than the number of data points [8]. In these cases, additional assumptions in the methods are necessary to make inference possible [8]. Additionally, some methods consider sedaDNA as binary occurrence (presence-absence) data, while others consider relative read abundance data [28]. These methods assume that relative read abundance is a proxy to organismal abundance or relative biomass, but there are many potential confounders that may affect this metric [29].

Understanding the relationships between these models and the circumstances under which they succeed or fail to infer species associations will be informative as the availability of sedaDNA increases. In particular, although we find that many of these methods do not perform well with realistic data and currently available sample sizes (see Results), we are hopeful that this will be informative as data availability increases. Additionally, one of the areas of promise presented by sedaDNA is the fact that we can potentially study the effects of taxa across kingdoms using the same samples [1], but in order to maximize this potential we will need to understand the performance of different methods across a broad range of ecological contexts.

Models to infer species associations have been tested against each other within the category of (J)SDMs [11, 30] and within the microbial network modeling literature [8, 27], but little is known about the relative performance of methods in the two categories. Often, these methods have been tested using data simulated under simple models that do not directly incorporate ecological processes [8, 11, 30]. Due to the complexity of real ecosystems, this may severely underestimate false inference rates of these models because the simulated data likely meets their assumptions much better than real data does. Therefore, it is important to also test methods under more challenging conditions that are based on ecological theory. To this end, we have developed a novel simulation model that uses ecological theory to simulate species abundances and simulates the sedaDNA sampling process. We compare this to a simpler simulation model that simulates sedaDNA read counts for multiple species without simulating ecological processes. We specify inter-species and species-environment interactions to create datasets for which the true interactions are known and apply different inference methods to the simulated data to test accuracy across a range of models, parameters, and datasets (Figure 1).

**Figure 1:**
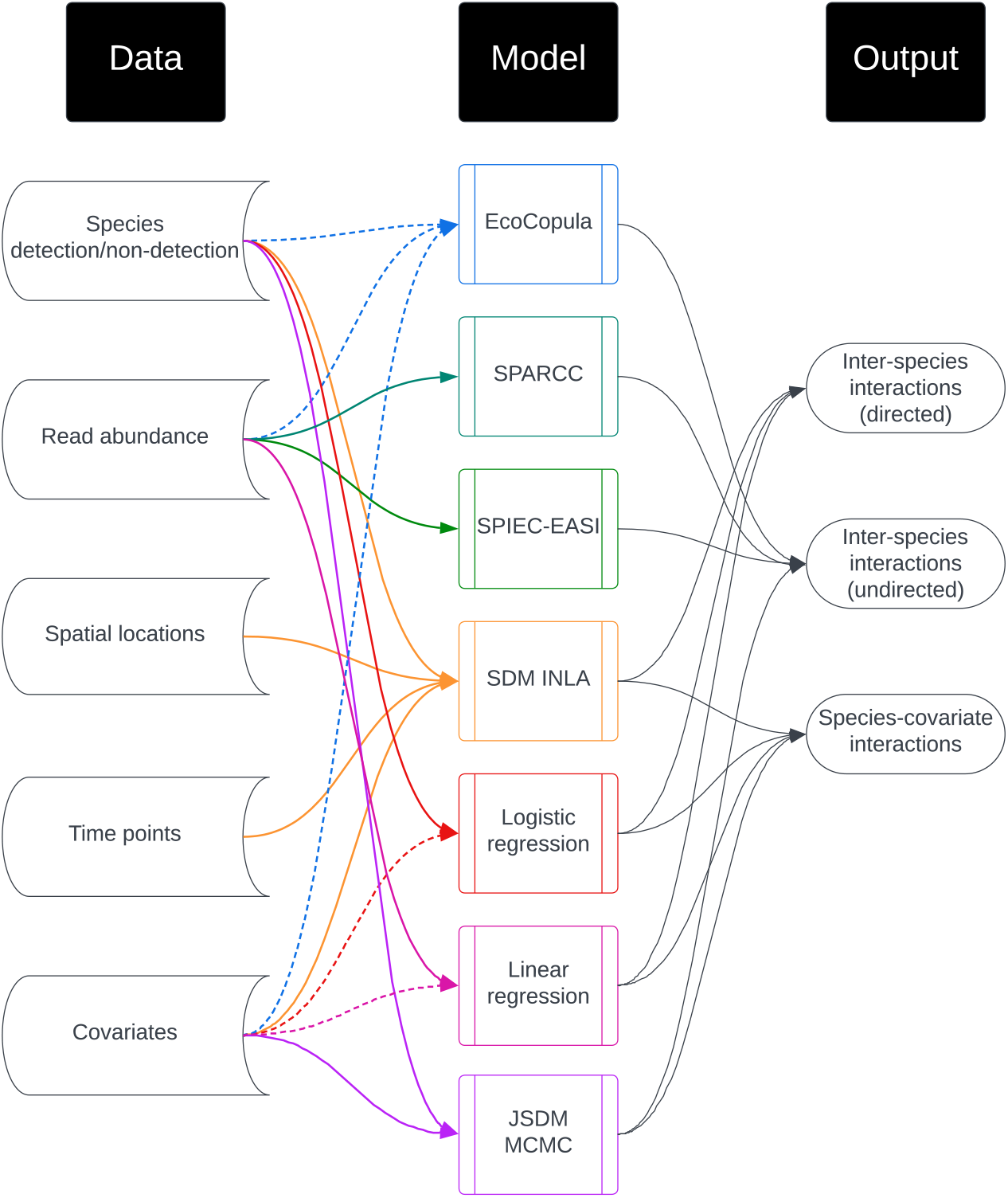
Inputs and outputs of inference methods tested. Dotted lines indicate optional inputs.

## Materials and Methods

### Simulation Models

Simulated data are generated under several different models. The first set of simulations uses a multivariate probit regression model modified to accommodate read counts, and is referred to here as *covariance matrix simulations*. In this model, species are simply associated via covariances in a latent Gaussian variable that is used to generate read abundances and presence-absence data. However, because we believe that ecological data can be much more complex than this model, we also simulated data under a more complex model that incorporates ecological theory, which we refer to as the *ecological simulation* model. We allowed parameters in this model to vary widely, but since we recognize that this may create unrealistic scenarios, replicates with a single set of parameters that generated more realistic data were also performed.

The *covariance matrix simulations* are designed to be as favorable as possible to the inference methods by violating their assumptions as little as possible. On the other hand, the *ecological simulations* introduce dynamics that violate many assumptions of the inference methods, so we expect that the methods will not perform as well under these conditions. The purpose of these models is to explore the robustness of these methods to a variety of realistic violations of their modeling assumptions.

### Notation

Throughout, *j* indicates species (used interchangeably with taxon, OTU, or ASV), ***s*** indicates a location in 2-dimensional space, and *t* indicates time. An omitted species index indicates the vector of values for all species. Bold symbols indicate vectors or matrices. Notation specifics can be found in Tables 1, 2, and 3.

**Table 1:** Variables and parameters in *covariance matrix simulation* model.

|  |  |
| --- | --- |
| $b$ | interacting block identifier |
| $J$ | total number of species |
| $J_b$ | number of species in interacting cluster $b$ |
| $F$ | number of latent variables for each block |
| $\Lambda_b$ | latent factor matrix for block $b$ |
| $\Gamma_b$ | intermediate covariance matrix before rescaling for block $b$ |
| $\Sigma_b$ | covariance matrix for block $b$ |
| $\Sigma$ | full covariance matrix for all species |
| $\mathbf{X}$ | matrix of covariate values for all sampling locations |
| $\beta$ | matrix of covariate effects for all species and covariates |
| $\mu^{(z)}$ | mean of $\mathbf{z}$ |
| $z_j(\mathbf{s}, t)$ | latent Gaussian variable used to define data (species $j$, location $\mathbf{s}$, time $t$) |
| $a_j(\mathbf{s}, t)$ | per-species read abundance data (species $j$, location $\mathbf{s}$, time $t$) |
| $y_j(\mathbf{s}, t)$ | binary species presence-absence data (species $j$, location $\mathbf{s}$, time $t$) |

**Table 2:** Variables in *ecological simulation* model. Variables change within a single simulation, in contrast to parameters, which are only chosen once per simulation.

|  |  |
| --- | --- |
| $\mathbf{s}$ | point in 2-dimensional space (unspecified units) |
| $t$ | point in time (unspecified units) |
| $j$ | taxon/species/ASV/OTU identifier |
| $n_j(\mathbf{s}, t)$ | number of individuals that deposited DNA |
| $a_j(\mathbf{s}, t)$ | sedaDNA read abundance data (species $j$, location $\mathbf{s}$, time $t$) |
| $y_j(\mathbf{s}, t)$ | binary species presence-absence data (species $j$, location $\mathbf{s}$, time $t$) |
| $N_j(\mathbf{s}, t)$ | abundance (species $j$, location $\mathbf{s}$, time $t$) |
| $K_j(\mathbf{s}, t)$ | carrying capacity of the environment (species $j$, location $\mathbf{s}$, time $t$) |
| $M_j(\mathbf{s}, t)$ | number of migrants (species $j$, (to) location $\mathbf{s}$, time $t$) |
| $\mathbf{x}^*(\mathbf{s}, t)$ | measured covariates |

**Table 3:** Parameters in *ecological simulation* model. Parameters change between simulation but are constant within a simulation.

|  |  |
| --- | --- |
| $J$ | number of species in the model |
| $P$ | number of covariates (environmental/abiotic factors) |
| $\beta$ | $J \times P$ matrix of covariate effects on the different species |
| $\alpha$ | $J \times J$ matrix of species effects on each other. Note: diagonal is 0 |
| $c_i$ | tuning parameters |
| $\lambda_j$ | migration rate for species $j$ |
| $r$ | intrinsic population growth rate |
| $\sigma$ | constant controlling amount of noise in population growth |
| $d$ | migration neighborhood radius for all species |
| $p_n$ | organism sampling rate |
| $\lambda_a$ | read sampling rate per organism |
| $\mathbf{x}(\mathbf{s}, t)$ | vector of environmental explanatory variables (covariates) |
| $\sigma_x$ | standard deviation for the covariate measurement noise |
| $R$ | detection threshold for presence-absence |
| $S$ | set of spatial locations |
| $V$ | scaling parameter for the spatial covariance function for $\mathbf{x}(\mathbf{s}, t)$ |
| $T_x$ | temporal period of the mean of for $\mathbf{x}(\mathbf{s}, t)$ |

### *Covariance matrix simulation* model

In one set of simulations, species interactions are encoded through the covariance matrix of a latent multivariate normal variable. The covariance matrix **Σ** for this set of simulations is defined as a *J* × *J* matrix with clusters of interacting and non-interacting species. In the clusters of interacting species, the inter-species covariances are defined using a latent factor model [31], such that all covariances are nonzero in a cluster but the correlation values vary as described below (empirical distribution of covariances in Appendix N).

Let *J*_*b*_ be the number of species in the interacting cluster, and let *F* be the number of latent variables. Then we define **Λ**_*b*_ as a *J*_*b*_ × *F* matrix with standard normally distributed components. For each cluster *b*, the entries in **Λ**_*b*_ are

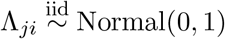

for all *j* ∈ {1, …, *J*_*b*_}, *i* ∈ {1, …, *F*}.

Now let **Γ**_*b*_ be the intermediate covariance matrix before rescaling. Define **Γ**_*b*_ as

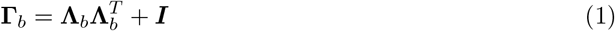

where ***I*** is the identity matrix with dimension *J*_*b*_ × *J*_*b*_. Then we re-scale **Γ**_*b*_ to a correlation matrix as follows:

Define

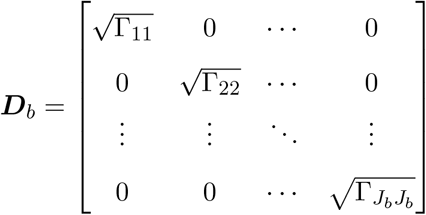

Finally, let the covariance matrix (also a correlation matrix in this case) for block *b* be defined as

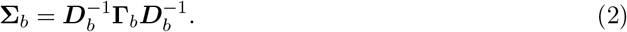

Now, for clusters *b*_1_, …, *b*_*n*_ we define the final covariance matrix **Σ** as a block matrix where 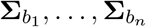 fill independent blocks on the diagonal and all other off-diagonal entries are 0. All entries on the diagonal are 1. Note that by defining **Σ** in this way, we have some species in clusters that are correlated to one another, and others which are statistically independent.

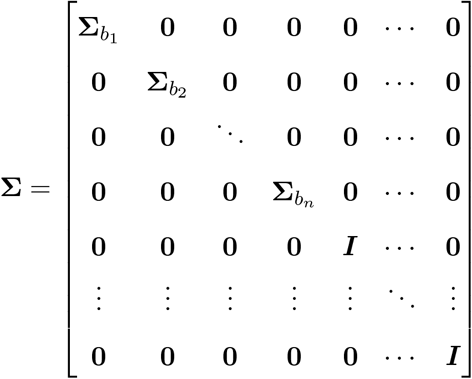

We define a set of clusters such that there are approximately the same number of interactions as species to maintain sparsity of interactions (as is assumed by some of the methods tested) [8]: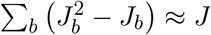.

We also include covariates, representing the background environment. We simulate covariates and their effects following Wilkinson, et al. (2019) [31]. We first simulate a matrix of four covariates plus an intercept, which vary over time and space. Call this matrix ***X***, which has dimension 5 × *N*, where *N* is the number of samples we are simulating (number of points in space-time, which for this model are statistically independent of each other). For the four non-intercept columns, we simulate matrix entries as 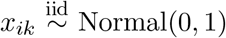 for *i* ∈ {1, …, *N*} and *k* ∈ {1, …, 4}. The intercept column is filled with 1’s. Now we simulate the effect of each covariate on each species, which we store in the matrix ***β***. This will be a *J* × 5 matrix. For some simulations we will fill ***β*** with 0’s, which denotes no effect of the environmental covariates on the species. For others, all entries of ***β*** will be independent and identically distributed standard normal variables. Now define ***µ***^(*z*)^ = ***βX*** as the *J* × *N* matrix of means for the latent multivariate normal variable. Each ***µ***^(*z*)^(***s***, *t*) will be a single column of this matrix, and will therefore be a vector of length *J*. Note that when the entries in ***β*** are nonzero, this introduces correlation structure between the species that is different from the correlations defined in **Σ**. For example, if by chance two species have similar responses to the environment, then their presence and read abundance will be positively correlated even if the corresponding entry in **Σ** is 0.

Now we simulate the latent variable ***z***, which will be used to produce both presence-absence and read abundance data. Let ***z***(***s***, *t*) be a length-*J* vector of latent multivariate normal variables which are correlated across species according to **Σ**.

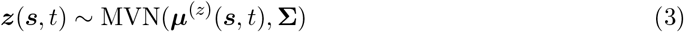

Now we will use these latent variables to generate presence-absence data and read abundance data.

Let ***a***(***s***, *t*) be the vector of simulated read abundances for all species at location ***s*** and time *t*. We then define the abundance as a Poisson random variable with mean proportional to the latent variable *z*_*j*_(***s***, *t*) if *z*_*j*_(***s***, *t*) is positive, or 0 otherwise. For all *j*, ***s***, *t*:

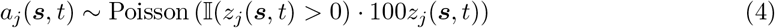

Define the presence-absence data ***y*** as 1 if *z*_*j*_(***s***, *t*) is positive or 0 otherwise. For all *j*, ***s***, *t*:

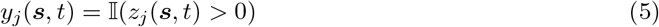

For the presence-absence data, this is a multivariate probit regression model. Due to the mean-zero truncated Gaussian, approximately half of the read counts are 0, which may be more or less realistic depending on the dataset. In this case, this is chosen to create optimal conditions for presence-absence data, as these methods lose power when percent presence is very low or high (Appendix M). Additionally, the latent variables are multiplied by 100 to put the mean of the Poisson distribution on the same order of magnitude as several example datasets, although we also recognize that this may vary between studies and depend on the level of classification of reads (e.g. OTU vs. ASV vs. species vs. family levels) [8, 20].

### *Ecological simulation* model

We constructed a simulation model which takes as input 1) inter-species and species-environment interactions, 2) environment covariates, and 3) simulation hyperparameters (e.g., life history traits, detection rates). The output of the simulation is sedaDNA read abundance and presence-absence data for each species at each point in time and space. Presence-absence data is created by setting a threshold and mapping the read abundance data to binary data.

This is a population-level model in which all individuals of the same species share the same dynamics and traits. It includes space explicitly, over which abiotic environmental covariates can vary in both space and time and an arbitrary number of species whose populations vary over space and time.

At each time step, we model migration and logistic population growth that depends on a time-varying carrying capacity. The carrying capacity at each time point and location is a function of abiotic covariates and the abundances of other species. We also model the detection process, including modeling DNA deposition, covariate measurement uncertainty, and varying numbers of sampled points (Figure 2).

**Figure 2:**
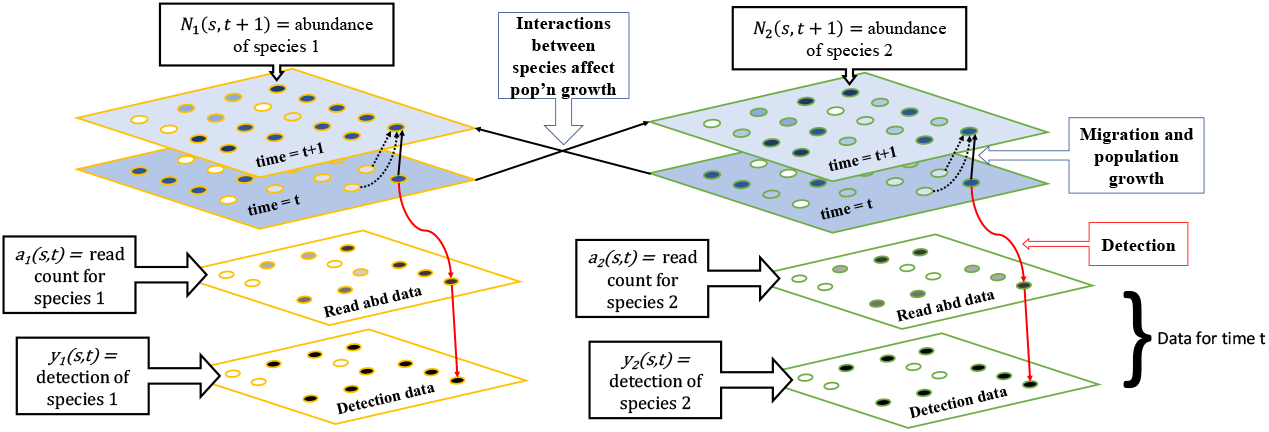
*Ecological simulation* diagram. At each time step, we model local migration and logistic population growth, which depends on the abundance of other species with specified interactions and a set of abiotic factors. We also model a sedaDNA detection process, resulting in two types of data: read abundance and presence-absence (detection) data. Arrows indicate the flow of information through the simulation. Dotted black arrows indicate migration, solid black arrows indicate population growth and species interactions, solid red arrows indicate data simulation.

Interactions are assumed to have an effect on carrying capacity. For example, competition between two species is represented through a lower carrying capacity for one species when the abundance of the other species is higher. Trophic interactions between species can also be represented in this way since a higher abundance of prey may increase the carrying capacity of the environment for the predator. Conversely, a higher abundance of a predator may decrease carrying capacity for its prey. Similarly, mutualistic relationships may be represented as a positive relationship between abundance of one species and carrying capacity of the other. Associations between species arise as an emergent property of these simulated interactions.

Note that variables (Table 2) change within a simulation, whereas parameters (Table 3) can be changed between simulations but are constants in a given simulation.

#### Mechanistic model of abundance

We model abundance as logistic growth in discrete time with noise and migration from a neighborhood of locations. Simulations were initialized with species abundances at time zero of *N*_*j*_(***s***, 0) = 10 for all species and locations, where *N*_*j*_(***s***, *t*) is the species abundance of species *j* in location ***s*** at time *t*. The first 100 time points are discarded before analysis.

The algorithm then proceeds by simulating the abundances as a two-step process as follows. At every time point *t >* 0:

Migration: At location ***s***, the number of new immigrants at time *t* for species *j* is *M*_*j*_(***s***, *t*), which is assumed to be Poisson distributed at a rate that depends on the number of individuals of species *j* that were in a neighborhood of location ***s*** at time *t*, and a species-specific migration rate *λ*_*j*_. Note that migrating individuals are not subtracted from the population they come from, which may be unrealistic for some scenarios but realistic for some others where the dispersal mechanism uses propagules rather than individual movement. The migration rate varies across species, with the actual value for each species drawn from a Gamma distribution parameterized with the mean across species set to *µ*_*M*_. The radius of the neighborhood, *d*, and mean migration rate are simulation parameters (allowed to vary between simulations but constant for a given simulation). The abundance in each location is then adjusted as follows:

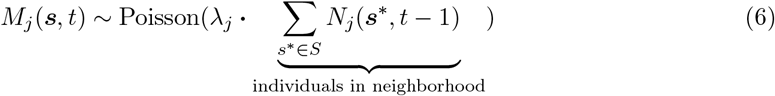

where *S* is the set of spatial locations within radius *d* of location ***s*** and *λ*_*j*_ ~ Gamma(shape = *µ*_*M*_ */*0.01, scale = 0.01). Then the migrating individuals are added to the population to form an intermediate population level, 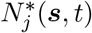.

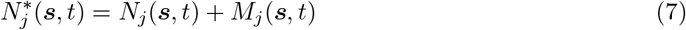

Population growth: The change in population size between time *t* and time *t* + 1, is modeled using discrete logistic growth with Gaussian noise and with a carrying capacity that depends on other biotic and abiotic factors. The growth rate of all species, *r*, and the noise of the process, scaled by *σ*, can be adjusted for each simulation but are constant within a simulation. We then assume that the local change in population size is

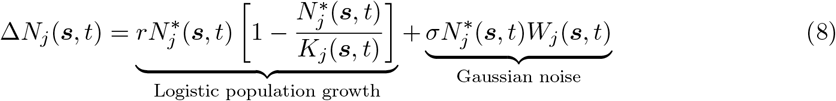

where *W*_*j*_(***s***, *t*) ~ Normal(0, 1).

The variance of the Gaussian noise term in the equation above scales with the population size as in ref. [32].

We model carrying capacity as a linear function of covariates and a non-linear function of other species abundance. The vector of carrying capacities for all species, ***K***(***s***, *t*), in the logistic growth equation depends on simulated abiotic environmental conditions, ***x***(***s***, *t*), and the abundance of the other species at time *t* − 1. The direction of the effect of each covariate and each species on other species is defined by matrices ***β*** and ***α. β*** is a *J* by *P* (number of covariates) matrix where each entry in the matrix indicates the sign and direction of influence of a particular covariate on a particular species. ***α*** is a *J* by *J* matrix where each entry in the matrix indicates the sign and direction of influence of a species on another species. In order to avoid inter-species effects going to infinity, the arctangent function is applied to species effects on each other. This function has a horizontal asymptote, creating an upper bound on the effect of one species on another, representing saturation of the inter-species effect. *c*_1_ and *c*_2_ are constants that control the relative strength of the effects of other species versus the abiotic environment on the growth rate of each species. Carrying capacities are truncated at 0 since they should not be negative.

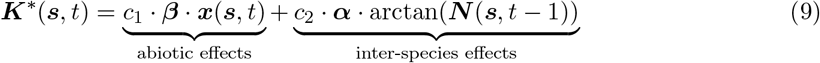

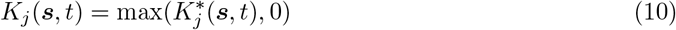

**Covariates (*ecological simulation*)** :

Covariates are simulated in two different ways for different *ecological simulation* sets (See Simulation Sets section of Methods).

Method 1: For the set-parameter simulations, covariates are simulated using Gaussian random walks through time. Covariates are therefore autocorrelated in time but not in space in this case. The vector of covariates for each time and location, ***x***(***s***, *t*), is an input to the population simulation model.

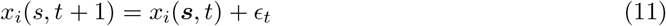

where *ϵ*_*t*_ ~ Normal(mean = 0, sd = 0.01).

Method 2: In the random-parameter simulations, covariates were simulated using a spatial Gaussian random field with an exponential covariance function, where the global mean varied in time according to a deterministic sinusoidal function. The period of this function, *T*_*x*_, and the spatial covariance parameter for the covariance function, *V*, are set uniformly at random within a set range for each simulation (See Table 4).

**Table 4:** Parameters in random simulation model. Parameters without a species index were set to be the same for all species, times, and locations in a given simulation.

| Parameter | Settings in random simulations |
| --- | --- |
| $J$ | 10 or 100 based on simulation set |
| $P$ | Uniform($\{3, 4, 5, \dots, J\}$) |
| $\beta$ ($J \times P$ matrix) | Uniform randomly selected $J$ entries set as 1 or -1, all others set as 0 |
| $\alpha$ ($J \times J$ matrix) | Uniform randomly selected $J$ entries set as 1 or -1, others set as 0 |
| $c_1$ | 200 |
| $c_2$ | Uniform(min = 100, max = 500) |
| $\lambda_j$ | Gamma(shape = $\mu_M / 0.01$, scale = 0.01) (For each species) |
| $\mu_M$ | Uniform(min = 0, max = 0.1) |
| $r$ | Uniform(min = 0.01, max = 1) |
| $\sigma$ | Uniform(min = 0, max = 0.2) |
| $d$ | Uniform(min = 12, max = 50) |
| $p_n$ | Uniform(min = 0.001, max = 0.02) |
| $\lambda_a$ | Uniform(min = 0.5, max = 100) |
| $\mathbf{x}(\mathbf{s}, t)$ | See covariate section |
| $\sigma_x$ | Uniform(min = 0, max = 0.2) |
| $R$ | Uniform(1, 2, 3, ..., 50) |

For each time point, all covariate values across space are drawn from a spatial random field with an exponential covariance function as follows:

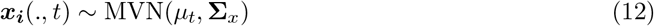

with

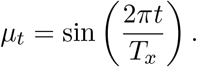

The mean value of the spatial random field (global mean of the covariate), vary in time according to a sine function. *T*_*x*_ is the period of the sine function in time, and it is drawn uniformly between 100 and 10,000 once per simulation:

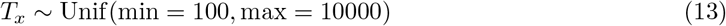

**Σ**_*x*_ is a spatial covariance matrix which is calculated as follows: for spatial points ***s***_2_ and ***s***_2_:

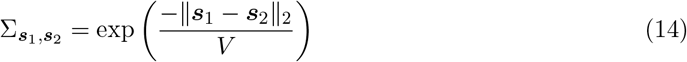

where ∥***s***_1_ − ***s***_2_∥_2_ is the Euclidean distance between the points and *V* is drawn once per simulation as:

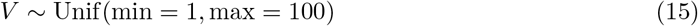

The covariance between two points decays exponentially as the distance increases, but it decays slower with increasing *V*.

#### sedaDNA read abundance and species detection model

We model sedaDNA by assuming that first, *n*_*j*_(***s***, *t*) individuals from the population are selected to deposit DNA. *n*_*j*_(***s***, *t*) is Binomially distributed with parameters *N*_*j*_(***s***, *t*) and *p*_*n*_, the individual sampling probability. The mean of this distribution is therefore proportional to the true species abundance, *N*_*j*_(***s***, *t*). This is equivalent to flipping a weighted coin with weight *p*_*n*_ for each individual present in the location to decide whether it deposits DNA. Then, the number of sedaDNA reads, *a*_*j*_(***s***, *t*), is Poisson distributed with a rate proportional to the number of DNA-depositing individuals, *n*_*j*_(***s***, *t*). The parameter *λ*_*a*_ dictates the rate at which each individual deposits DNA. In other words:

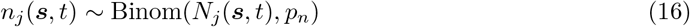

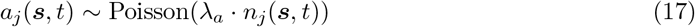

Presence-absence data, *y*_*j*_(***s***, *t*), is created by truncating the read abundances at some threshold *R*.

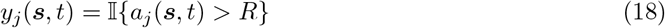

In the ecological model simulations, under many parameter settings, some species went extinct quickly, so species were filtered for at least 10% presence in the dataset. Therefore, the number of species actually analyzed is less than the number of species simulated in many cases. However, if the number of species after filtering was less than half the number originally simulated, the data set was discarded.

### Covariate measurement uncertainty

Covariates are measured with Gaussian noise with variance *σ*_*x*_:

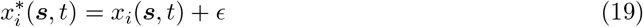

and *ϵ* ~ Normal(0, *σ*_*x*_).

### Simulation Sets

Simulation sets were designed with three main axes of stratification: simulation design (*covariance matrix simulation* and *ecological simulation*), number of species (~ 10 and ~ 100), and sample size (100, 250, and 10,000). The different simulation designs and three sample sizes represent different levels of realism. Ecological data is inherently complex, so all methods must make assumptions that are potentially violated by real data. However, methods may be differentially robust to these violations. If a method performs poorly on realistic data, this may be attributable to several causes. The assumptions of the method and the actual data generating process may be too different for the method to perform well, or the amount of data may be too small. We can differentiate between these different types of problems by examining performance at a realistic sample size (100 samples), a large but potentially feasible sample size (250 samples), and an unrealistically large sample size (10,000 samples). If a method is statistically consistent, when its assumptions are met it should converge to the true solution as the sample size gets large. Therefore we can attribute large errors at a large sample size to violations of modeling assumptions.

#### *Covariance matrix simulations* without covariate effects

We simulated 100 datasets with 10 species and 100 datasets with 100 species. For the datasets with 10 species, there was one interacting block with 4 species for a total of 12 one-way interactions. In the datasets with 100 species, there were 5 interacting blocks, each with 5 species, for a total of 100 one-way interactions. For this simulation, the covariate effects ***β*** were all set to 0. Although the covariates did not affect the species data, environmental variables themselves were still simulated. Although time and space are not included in this simulation algorithm, spatial and temporal labels were included in the final dataset to allow all methods to be tested. Each simulation was subsampled to 100 samples, 250 samples, and 10,000 samples for analysis.

#### *Covariance matrix simulations* with covariate effects

We simulated 100 datasets with 10 species and 100 datasets with 100 species. Everything was the same as the set without covariate effects except that the covariate effects ***β*** were each set to be independent standard normal variables. In other words, for all 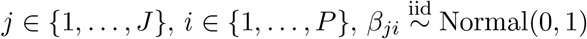.

#### Random-parameter *ecological simulations*

We ran 100 simulations under the ecological model described above with all parameters set uniformly at random in what we assessed to be a reasonable range (ranges and distributions specified in Table 4). In each simulation, 100 spatial locations arranged in a 10 by 10 grid were simulated for 10,000 time points, but the full simulated dataset was not analyzed. We simulated 100 replicates each with 10 species and 100 species. We randomly re-sampled each dataset to 100 samples, 250 samples, and 10,000 samples for analysis.

#### Set-parameter *ecological simulations*

As these random-parameter simulations often lead to scenarios with poor performance of most methods (see Results), we also used a model with fixed parameters for which we expect somewhat better performance as many sources of noise are set to a low level. For example, measurement noise in the covariates was set at 0, the detection rate was relatively high, and noise in population growth was set low. Specific parameter values are described in Table 5. Using these parameter settings, we simulated 100 replicates each with 10 species and 100 species. We randomly subsample each dataset to 100, 250, and 10,000 samples.

**Table 5:** Parameters in set-parameters simulation. Parameters without a species index were set only once per simulation for all species, times, and locations.

| Parameter | Setting in set-parameter simulations |
| --- | --- |
| $J$ | 10 or 100 based on simulation set |
| $P$ | $J$ |
| $\beta$ ($J \times P = J$ matrix) | $I_J$ (Identity matrix) |
| $\alpha$ ($J \times J$ matrix) | Randomly selected $J$ entries set as -1 or 1, others set as 0 |
| $c_1$ | 200 |
| $c_2$ | 300 |
| $\lambda_j$ | Gamma(shape = $\mu_M/0.01$, scale = 0.01) |
| $\mu_M$ | 0.01 |
| $r$ | 0.05 |
| $\sigma$ | 0.005 |
| $d$ | 16 (only direct neighbors) |
| $p_n$ | 0.01 |
| $\lambda_a$ | 1 |
| $x(\mathbf{s}, t)$ | See covariate section |
| $\sigma_x$ | 0 |
| $R$ | 5 |

### Testing inference models

All methods were used with default settings in the papers cited, with the exception of threshold adjustment to make receiver operating characteristic (ROC) curves. All methods that accept but do not require covariates (logistic and linear regression, SDM-INLA, EcoCopula) were tested both with and without covariates included in the analysis, regardless of whether the covariates had an effect on the simulated species data (Figure 1).

#### Logistic Regression

Logistic regression was performed using the `glm` function from the `stats` package version 4.4.0 [33]. Separate regressions were run for each species as a function of all other species (and covariates for runs with covariates). Interactions were then considered significant based on a p-value threshold. For reported false discovery rates at a single p-value threshold, the threshold was chosen according to the Benjamini-Hochberg procedure with expected false discovery rate of 0.05 [34]). For ROC curves, the threshold was varied from 0 to 1. Presence-absence data (and covariates, where applicable) were used as input to the model.

#### Linear Regression

Linear regression was performed using the `lm` function from the `stats` package version 4.4.0 [33]. Analyses thereafter proceeded as described for logistic regression. Read abundance data (and covariates, where applicable) were used as input to the model.

#### JSDM-MCMC

JSDM-MCMC was performed as in Pollock, et al., 2014 [14] using JAGS version 4.3.2 [35] and package R2jags version 0.7-1 [36]. It decomposed the species co-occurrence patterns into components describing shared environmental responses and residual patterns of co-occurrence including species interactions. For each species pair, it returned a set of MCMC samples of the association parameter, and significance was determined based on whether a q-percent credible interval contained 0. To calculate false discovery rate at a single threshold, the chosen q-percent was 95%. For ROC curves, the q-percent was varied from 0 to 100%. The model was run with covariates and with presence-absence data as the input. Convergence of the MCMC algorithm was evaluated by examining trace plots of multiple chains and using Gelman-Rubin statistics in the R package Coda [37, 38] (Appendix C).

#### SDM-INLA

For SDM-INLA, analysis was performed as in Wang, et al., 2021 [12] using INLA version 24.2.9 [39]. For model selection, WAIC was used to choose between four models: 1) using all other species and covariates as predictors, 2) using all other species as predictors without environmental covariates, 3) only environment covariates as predictors without other species’ data, and 4) no predictors. 95% posterior credible intervals were used to determine associations. Following Wang, et al. (2021), all models also included a spatiotemporal effect [12]. For ROC curves, no model selection was used (models (1) and (2) form two separate curves) and the cutoff for the quantile of the posterior credible interval was varied. Presence-absence data, covariates (where applicable), time points, and locations were used as input to the model.

JSDM-MCMC and SDM-INLA methods were not used in the simulations with a larger numbers of species because it took a prohibitively long time for the methods to run on the large number of datasets simulated here (Appendix A). Additionally, for SDM-INLA, the method often failed, but failure was not consistently repeatable, even using the same dataset. This could be avoided for individual datasets by re-running multiple times or optimizing settings specifically for individual datasets, but doing this for every dataset was not practical for this study.

#### SPIEC-EASI

SPIEC-EASI was run using the R package SpiecEasi version 1.1.2 [40]. All parameters were set to default, and both the `mb` version and the `glasso` version of the model were run for comparison. The input to the model was simulated read abundances. For ROC curves, the regularization parameter, `lambda`, was adjusted to 100 different values automatically by the SpiecEasi package (`nLambda = 100`) and then results were averaged across simulations for the ordered `lambda` values. The `lambda` values were ordered but not necessarily the same values across runs of the simulation, since by default the model automatically selects the actual values. For the model-selected results (reported false discovery rates), a specific `lambda` value was selected through the default calibration procedure.

#### SparCC

The SpiecEasi package also implements the model SparCC, which was originally published by Friedman and Alm (2012) [21]. This model, as implemented in SpiecEasi, was also tested on simulated read counts. For ROC curves, the threshold for estimated covariance, which is used to call interactions, was adjusted. For the false discovery rate after model selection, a pseudo p-value was generated using the bootstrapping procedure described by Friedman and Alm (2012) and implemented in SpiecEasi [21, 40]. The cutoff for this pseudo p-value was set at 0.05.

#### EcoCopula

EcoCopula was run using the R package EcoCopula version 1.0.2 [9]. The model was run with and without covariates. This method, unlike the others, can take flexible types of input data. Therefore, we tested it using the mode where it takes binary (presence-absence) data and where it takes count (read abundance) data. By default, the method selects 100 values to test for the regularization parameter, `lambda`. Then it chooses one using BIC. Default selection procedure for `lambda` was used for FDR results. ROC curves were produced by varying the `lambda` values (selected from those chosen by the default model) and averaging resulting true and false positive rates across simulations. The `lambda` values were ordered but not necessarily the same values across runs of the simulation. The `lambda` parameter was also set at 0 to complete the ROC curve, although this was not among the values selected by the model by default.

### Counting Mistakes

The performance of these models was evaluated in several ways in order to get a more complete picture of their performance. First, we evaluated the success of the methods at detecting direct, causal interactions. Since this type of data is generally observational, and therefore can (and likely does) have significant confounding variables, we do not claim that any of these methods would detect direct, causal relationships consistently in real data. However, we believe that it is still useful to examine whether they are able to detect causal relationships when all variables are observed. For this metric, sign of interactions was ignored in the calculation of false positive and negative rates. If an interaction exists and one was inferred, this was counted as correct regardless of the interaction being positive or negative. Interactions were considered as directed, so A influencing B does not imply B influencing A. Therefore, for example, false positive count was calculated as the number of times where an interaction was inferred from A to B, but no interaction exists from A to B, regardless of the sign being positive or negative (Figure 3).

**Figure 3:**
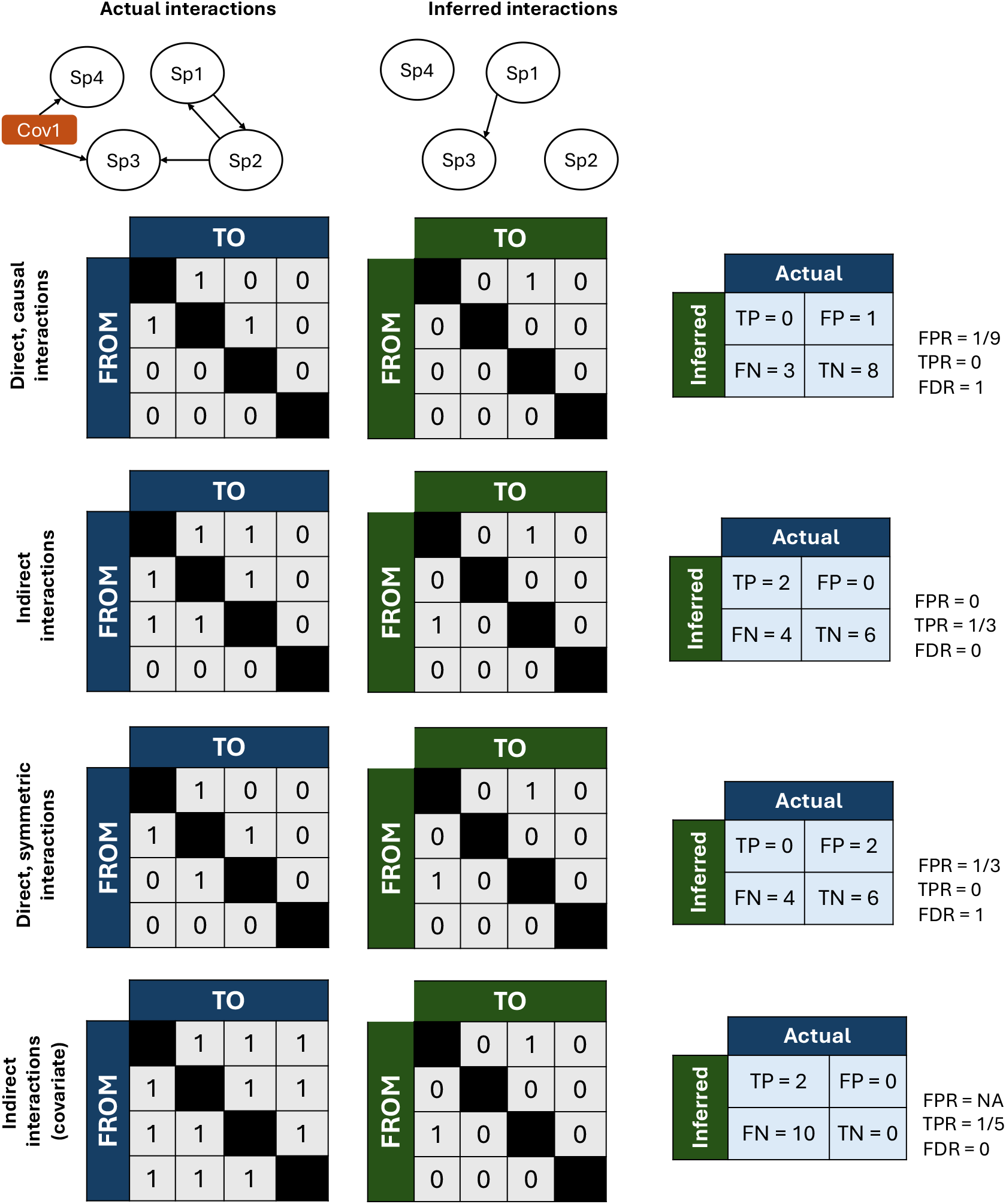
Schematic explaining the calculations of false positive rate (FPR), true positive rate (TPR), and false discovery rate (FDR), when interactions are considered direct (symmetric or asymmetric) versus indirect (with or without considering shared covariate interactions).

Second, we evaluated the success of the different methods at detecting whether there are any interactions between two species, whether they are direct or indirect through other species (indirect interactions). For this metric, we considered an interaction to exist if the two species are connected by interactions, regardless of the direction of the interactions. We likewise consider an interaction to be inferred between two species if any inferred associations connect them (Figure 3).

Third, we evaluate the success of the models at detecting direct associations between species without direction (direct, symmetric interactions). Some methods have the theoretical potential to infer an interaction in one direction but not the other (SDM-INLA, JSDM-MCMC, logistic regression, linear regression), although this should not happen often since they are all correlative methods, while other methods always infer symmetric interactions (SPIEC-EASI, SparCC, EcoCopula). Using this metric, if an interaction exists from A to B, then we automatically assume one exists from B to A. Likewise with the inferred associations, we assume that an inferred association in one direction implies one in the other direction.

Fourth, we consider indirect interactions to include interactions through covariates (indirect interactions, covariate). For example, if a covariate affects two species, these species are considered to interact. However, since interactions between covariates and species are not inferred, the inferred interactions were defined based on whether two species are connected by interactions between species, without regard for covariates. This metric did not cause different results than the indirect interactions metric without considering covariates for the *ecological simulations* and is not an informative metric for the *covariance matrix simulations* since all species are connected by covariates (See results), but we include it here for completeness.

False discovery rate was calculated as false-positive-count/total-inferred-associations, or the probability that an inferred interaction was incorrect. This is distinct from false positive rate, which was calculated as false-positive-count/total-actual-interactions. These metrics can differ enormously, especially when there is severe class imbalance, which is the case here with many more pairs with no actual interaction compared to the number of actual interactions (sparsity of the interaction matrix). Here, we present the false discovery rate after model selection, and the false positive rate at many thresholds as part of ROC curves.

### Analysis of effect of individual simulation parameters on predictive performance

A random forest model was used to predict false discovery rate for linear and logistic regression using the simulation parameters that were set at random in the simulations. Random forest models were run in R using Ranger version 0.16.0 [41]. The model was trained on 1000 simulations, with 10 species each and with 100 or 10,000 samples per simulation. Linear and logistic regression were both tested, using no covariates for either model, and with corrected p-values (Benjamini-Hochberg correction at a 0.05 false discovery control level). FDR was evaluated for direct/symmetric interactions only.

The following formula was used for random forest analysis (See Table 3 for definitions and Table 4 for ranges of parameter settings):

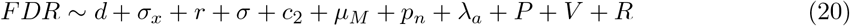

Variable importance in the random forest prediction was evaluated using permutation importance (`importance = “permutation”` in Ranger model). The random forest predictive performance was evaluated using root mean square error (RMSE) for an independent test set of 100 simulations. The RMSE of the random forest was compared to a naive predictor to evaluate how much predictive power the simulation parameters have for the resulting FDR. The naive predictor was the median FDR value for the dataset.

## Code

Simulations and SDM-INLA analyses were run in R version 4.3. All other analysis was run in R version 4.4 [33].

Code is available at https://github.com/Fiona-MC/eDNA-sims-pub.

## Results

Several methods that test for associations between different taxa (used interchangeably with species, OTU, or ASV) were tested for their effectiveness at detecting these associations. Data was simulated using two models: a mechanistic *ecological simulation* model and a simpler *covariance matrix simulation* model. The *ecological simulation* model was designed to be as realistic as possible, and since the complexity of ecological data necessitates making assumptions that real data may not follow, this means that this data violates many of the assumptions of the inference methods. On the other hand, the *covariance matrix simulations* are less realistic but also violate fewer of the assumptions of the inference methods, so we expect performance on this data to be better. These different simulation types examine robustness of the methods to differing levels of violations of modeling assumptions.

### False discovery rates are high for most inference models and simulations when assumptions are violated

For the *ecological simulations*, false discovery rate (FDR) for direct, causal interactions (Figure 3) in all cases was over 50% for 100 samples (Figure 4). For 250 samples, the same general trends apply, though the FDRs are lower in general, as expected, with the lowest being 38%. We also include simulations with 10,000 samples to see which mistakes are caused by an insufficient amount of data. For 10,000 samples in the *ecological simulations*, the lowest false discovery rate was still 50% (Figure 4). The only simulation for which the false discovery rates are generally low is the *covariance matrix simulation* with no covariate effects, which is the simulation that minimally violates assumptions of the methods.

**Figure 4:**
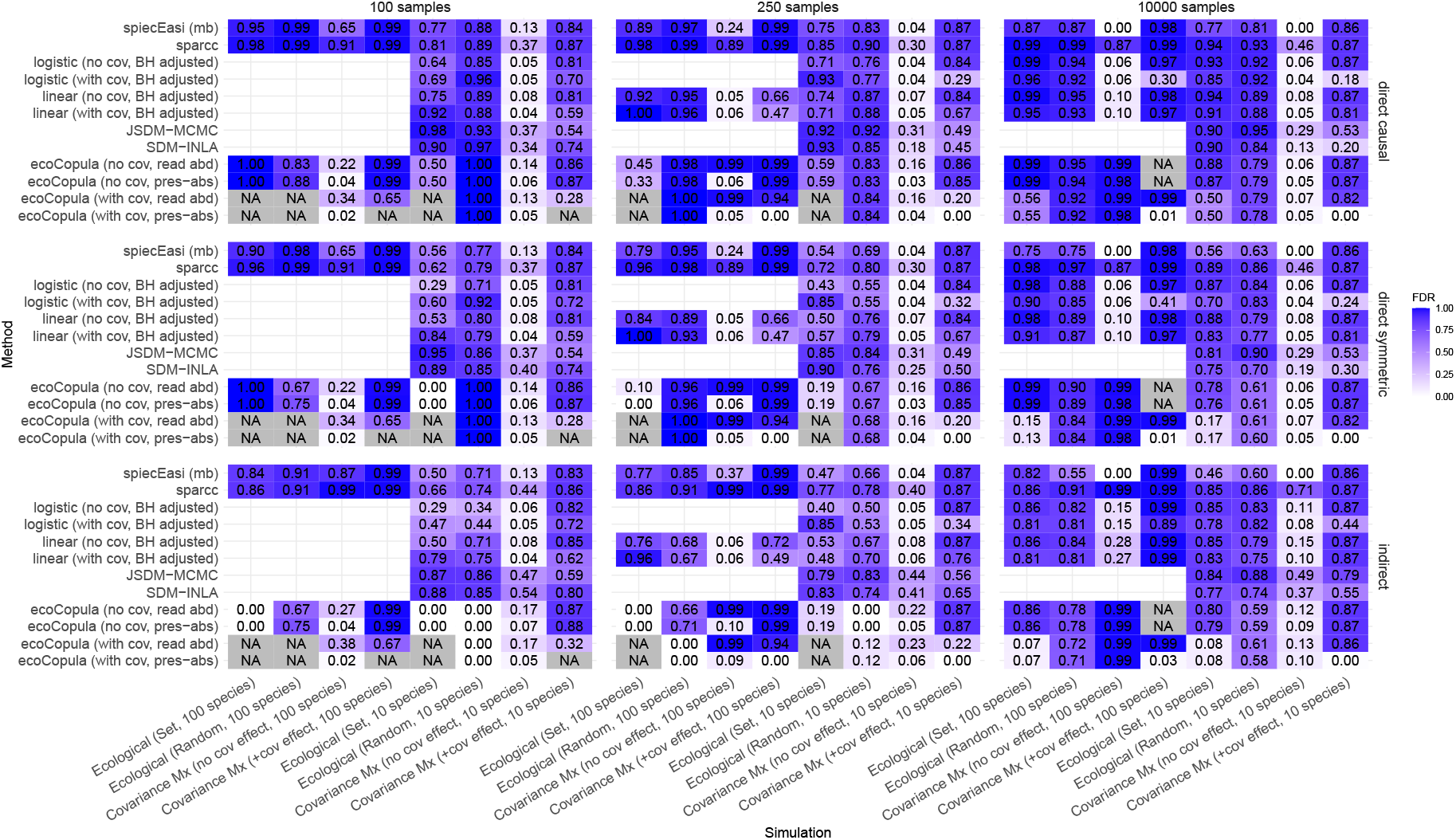
False discovery rates of direct/causal interactions, direct/symmetric interactions, and indirect interactions with 100, 250, and 10,000 samples. False discovery rate is defined as the number of false positive detections of species interactions divided by the total number of inferred interactions, with interactions defined in three ways. NA indicates that no interactions were inferred, and therefore false discovery rate is undefined. Where there is an option, cov refers to whether covariates were used as predictors in the inference methods or as environmental effects in the simulation. Similarly, where adjustable, `pres-abs` and `read abd` refer to presence-absence data versus read abundance data inputs respectively.

Some of the mistakes are caused by including the direction of the inferred associations in counting mistakes. For the *ecological simulation* sets, the false discovery rate decreases when interactions are considered symmetric, and for the *covariance matrix simulation* sets they are approximately the same. Interactions in ecology are often stronger in one direction than the other (i.e. species A affects species B but species B has little affect on species A), so we wanted to examine the effectiveness of these methods at detecting directed versus undirected interactions. In the *ecological simulation* model, some species have a unidirectional influence on other species, whereas in the *covariance matrix simulation* model, all inter-species effects are bidirectional and the species are organized in clusters so there is little difference between the three metrics shown. SDM-INLA, JSDM-MCMC, and the regression methods can in theory predict asymmetric associations, but all other methods predict associations symmetrically. These associations are nonetheless often interpreted as potentially having causal meaning. Therefore, we have included the direct, causal interactions to illustrate one of many reasons that these associations should not be interpreted as directional or causal without additional information.

False discovery rates using the metric of indirect interactions (Figure 3) vary dramatically between methods. For some methods and simulations, all inferred interactions are correct (FDR = 0), while for others, the false discovery rate is as high as 0.99 (nearly all inferred interactions incorrect) (Figure 4). EcoCopula and SPIEC-EASI claim to be able to avoid inferring indirect interactions by estimating conditional dependence of species presences (read abundances) given all other species presences (read abundances) [8, 9]. However, here we see that the false discovery rate for SPIEC-EASI is either comparable or goes down when interactions are considered indirect (Figure 4). Notably, this remains true when the model is simplified to the *alternative covariance matrix simulation*, which has a different simulation mechanism specified in Appendix G, including having interactions which are not organized in clusters (Appendix G). For EcoCopula, the story is less clear between considering interactions as direct but symmetric versus indirect. There are many cases where indirect interactions are inferred much better. In fact, using similar data, the number of interactions inferred after calibration varies considerably. It is possible that for a single dataset, this problem could be mitigated by fine-tuning the parameters based on the specifics of the data.

There are a few outliers in the FDR results for 100 samples that may be caused by very low overall rates of inferred interactions. For example, the 0% FDR for EcoCopula in the direct symmetric interactions case was caused by only 4 interactions that were all inferred correctly in one dataset (out of 100) (Figure 4). In all other datasets, no interactions were inferred. Similarly, for EcoCopula with no covariates for the set of simulations with 100 species, only 2 interactions were inferred in total, but both were incorrect, resulting in an FDR of 100% (Figure 4; Appendix B). When such low numbers of total interactions are inferred, the estimates of FDR may rely on just a few unusual cases and therefore may have high variance.

For the *covariance matrix simulations*, the three interaction metrics shown are quite similar. Any differences arises from transforming the inferred interactions to be symmetric or indirect, since the ground truth interactions are the same for all three. Whether interactions through the environment are considered correct or incorrect has almost no effect on FDR for *ecological simulations* (Appendix E). For the *covariance matrix simulations*, all species are connected through covariates and therefore this metric is uninformative.

### Model calibration (model selection or p-value cutoff) dramatically affects performance

False discovery rates are calculated using the default calibration for each method, but these calibration methods (model selection or choosing a p-value cutoff) vary between methods, which can cause large differences in the number of inferred interactions and the FDR. ROC curves do not depend on calibration, though the calibrated value is shown on the curves as the larger dot (Figure 5, 6, 7, 8). The number of total inferred interactions is as low as 0 for some (seen as NA in Figure 4), and up to over 9000 inferred interactions per simulation for others (Appendix B). In many cases, ROC curves look relatively good but the FDR is very high, indicating that the underlying model is able to discriminate between interacting and non-interacting pairs of species, but the calibration method is selecting a point on the ROC curve that results in a high FDR.

**Figure 5:**
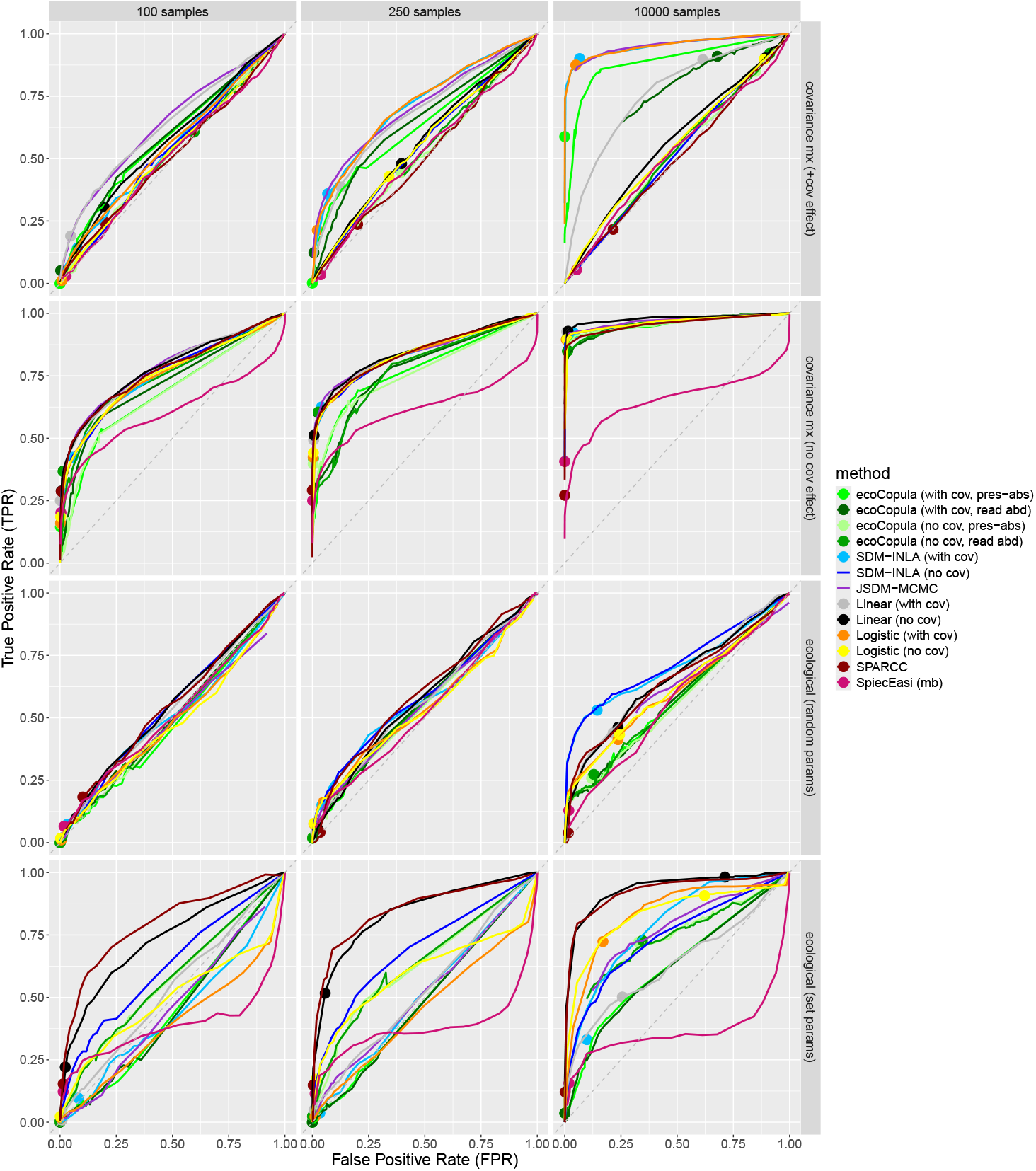
ROC curves for inference of direct, symmetric associations between 10 simulated species (5-10 species actually present). `cov` refers to whether covariates were included in the model, but in all cases, species interactions were inferred. `read abd` and `pres-abs` are specified for EcoCopula because this method allows for different inputs (presence-absence data or read abundance data). Solid points are the point chosen by the default model selection of each method.

**Figure 6:**
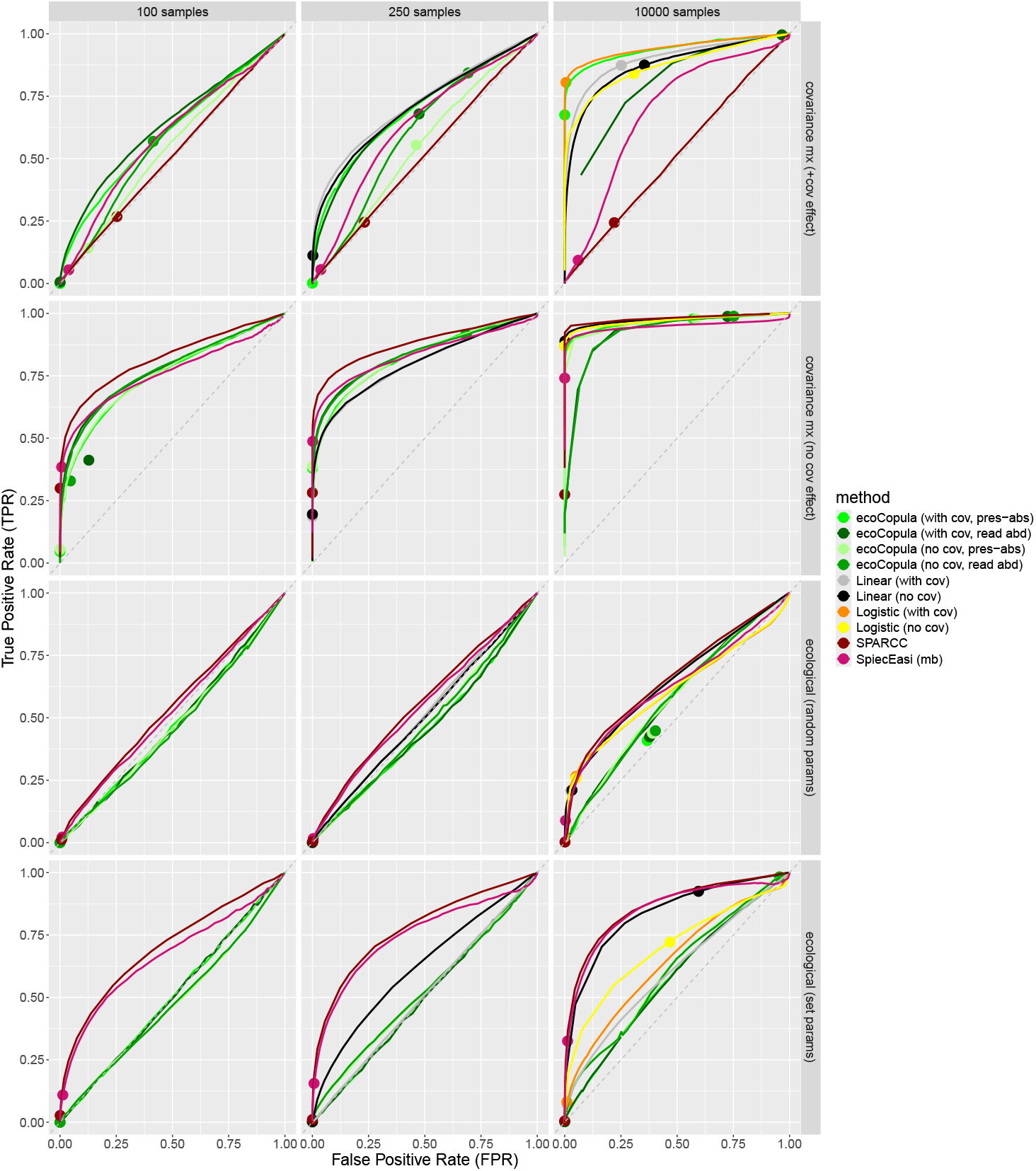
ROC curves for inference of direct, symmetric associations between 100 simulated species (50-100 species actually present). `cov` refers to whether covariates were included in the model, but in all cases, species interactions were inferred. `read abd` and `pres-abs` are specified for EcoCopula because this method allows for different inputs (presence-absence data or read abundance data). Solid points are the point chosen by the default model selection of each method.

**Figure 7:**
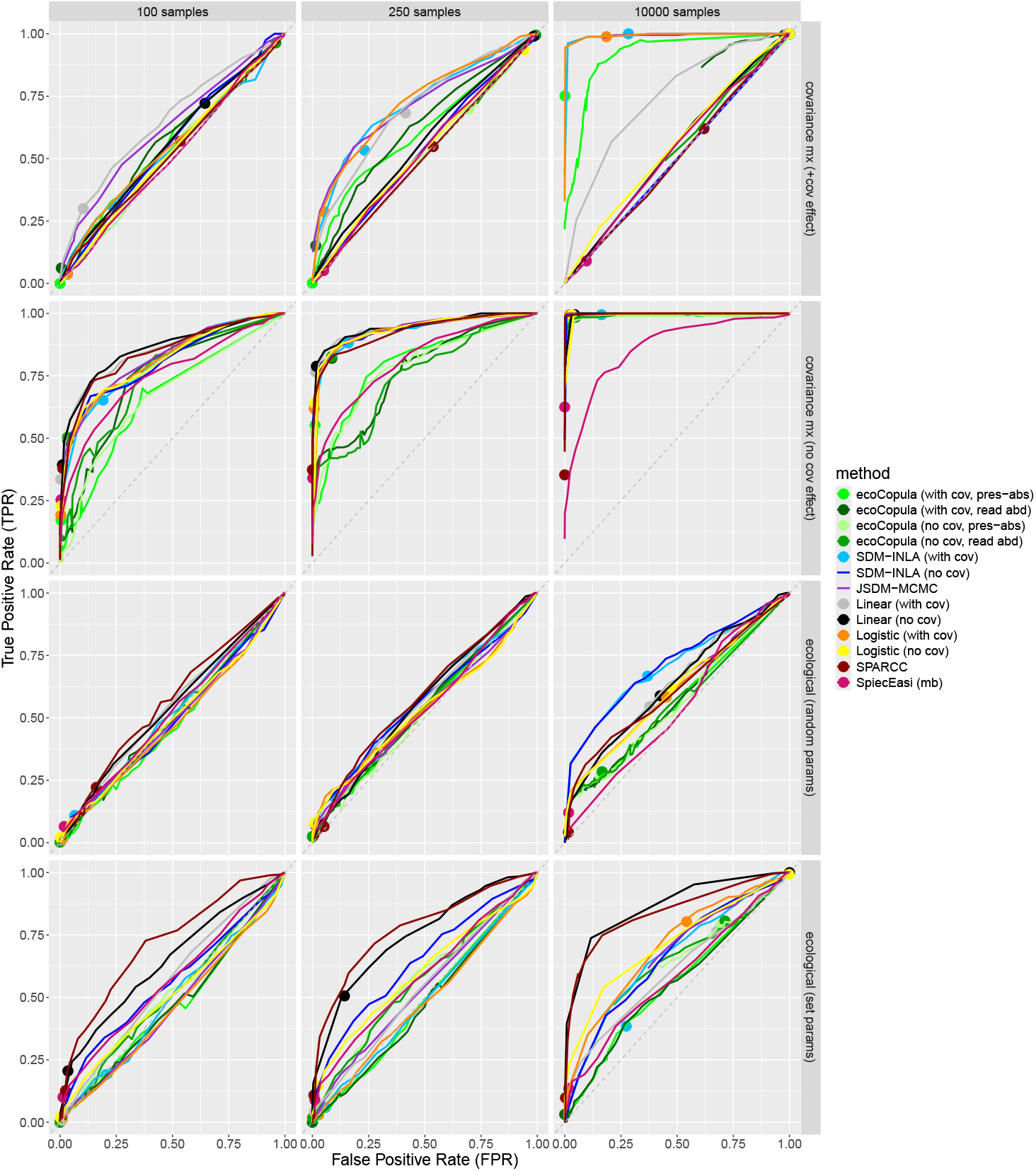
ROC curves for inference of indirect associations between 10 simulated species (5-10 species actually present). `cov` refers to whether covariates were included in the model, but in all cases, species interactions were inferred. `read abd` and `pres-abs` are specified for EcoCopula because this method allows for different inputs (presence-absence data or read abundance data). Solid points are the point chosen by the default model selection of each method.

**Figure 8:**
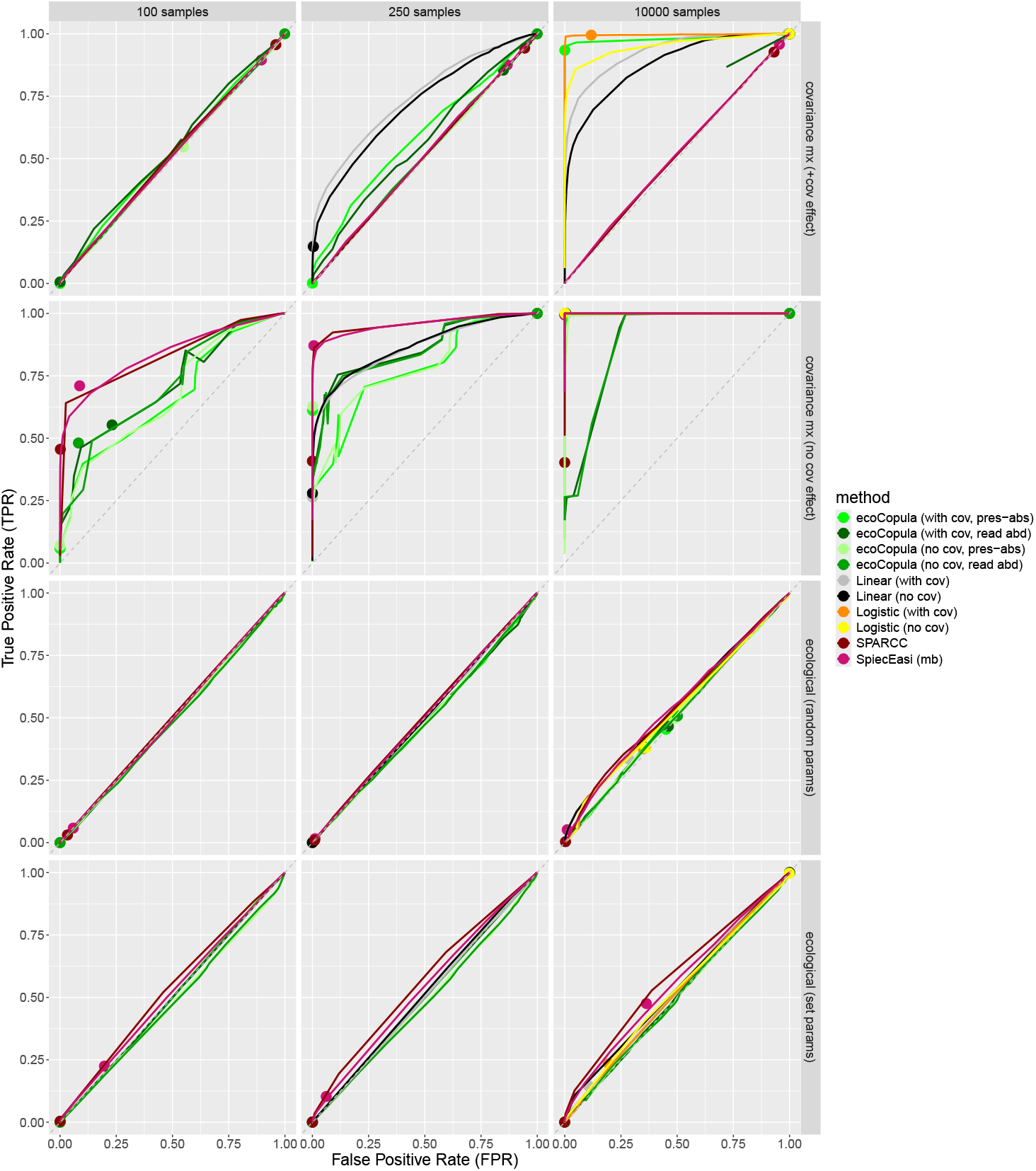
ROC curves for inference of indirect associations between 100 simulated species (50-100 species actually present). `cov` refers to whether covariates were included in the model, but in all cases, species interactions were inferred. `read abd` and `pres-abs` are specified for EcoCopula because this method allows for different inputs (presence-absence data or read abundance data). Solid points are the point chosen by the default model selection of each method.

As expected for lower amounts of data, when only 100 points were sampled many methods often inferred very few interactions. On the other hand, with 10,000 samples, hundreds or thousands of total interactions were inferred across the 100 simulations per dataset (Appendix B). One exception was SPIEC-EASI, which did not consistently infer fewer interactions when fewer points were sampled. In additional tests with as few as 10 samples, SPIEC-EASI continues to infer large numbers of interactions (Appendix D), but with so few samples it is unlikely that there is enough information in the data to infer this, indicating potential problems with calibration. Also, EcoCopula sometimes inferred drastically different numbers of interactions based on whether presence-absence or read abundance data was used as input. Interestingly, for some of these cases the ROC curves do not differ significantly, indicating a problem with calibration rather than the underlying model. For example, for the *covariance matrix simulation* with no covariates and 100 species, over 6000 direct associations per simulation were inferred for read abundance data (with and without covariates) and around 40 direct associations per simulation with presence-absence data (with and without covariates). In general we would expect that calibrated values would be somewhere on the ROC curve line. However, we see that this is not always the case for EcoCopula. The ROC curves are produced by varying the regularization parameter, but we find that manually setting the value of this regularization parameter sometimes has a different result than allowing the model to select the value using BIC, even when we attempt to manually select the same value as the optimal value that the model selects. Additionally, curves are sometimes not concave due to the effect of averaging across runs while changing the regularization parameter. We therefore recommend careful examination of sensitivity to calibration metrics and parameters in these methods.

We also examined different calibration methods for the regression methods, including no false discovery rate correction (least conservative), Benjamini-Hochberg adjustment, and Bonferroni adjustment (most conservative). As expected, more interactions were inferred and false discovery rates were generally higher with the less conservative methods (Appendix O). All of the results shown for regression use the Benjamini-Hochberg adjustment, with the results for other correction methods shown in Appendix O. For methods like SDM-INLA, the model is run separately for each species but no FDR correction is applied (instead, the model is selected using WAIC), which may contribute to high false discovery rates.

### High numbers of samples are needed to infer associations between species in more realistic scenarios

Increasing sample size is expected to improve the performance of statistical methods in most cases, although model misspecification can result in incorrect inference even as sample size gets very large [42]. Here, we have tested each model at a relatively low sample size, which we believe is realistic for sedaDNA studies at this time (100 samples) and a larger but still potentially feasible sample size (250 samples). We have also tested all models with 10,000 samples, which is a higher number of samples than would currently be reasonable in sedaDNA studies. However, it is useful to examine performance at a very high sample size since large errors can then be attributed to insufficient robustness rather than insufficient data. As expected, we find that model performance is better for most methods and scenarios using a higher sample size, although FDR remains high in many cases (Figure 4).

Using only 100 samples, on average for random parameter settings in the *ecological simulation* model and for the *covariance matrix simulations* with covariates, no method performs better than a random classifier (defined as choosing to infer interactions completely at random, which is expected to follow the diagonal on a ROC curve plot) (Figure 5, 6, 7, 8). The only set of simulations where the methods all consistently perform better than random using this small sample size was the simplest set of simulations where species have no response to environmental covariates, which is the least realistic scenario. For the *ecological simulation* with set parameters, some methods perform better than random and some still perform poorly with low sample sizes.

With model selection, it is often the case that very few interactions were inferred with only 100 samples, which indicates low power to detect these interactions if the method is calibrated correctly (Appendix B). When interactions are inferred, they are most often incorrect for *ecological simulations* and *covariance matrix simulations* with covariate responses (Figure 4). Therefore, we find it to be a good sign when methods consistently infer very few interactions, since no method is able to detect the correct set of interactions. As expected, interactions are inferred more often with a higher number of samples and are more often correct (Figure 4, 5, 6). However, there is still a high chance that inferred direct interactions are incorrect under many scenarios (Figure 4). Using higher sample sizes, the false discovery rate is only consistently low in very favorable scenarios (*covariance matrix simulation* without covariate effects; note exception of set-parameters with covariates under the EcoCopula model, though this relies on very few inferred interactions, and a few exceptions for the *covariance matrix simulations* with covariates) (Figure 4).

The level of correlation induced by species interactions in real data, which is expected to influence the necessary number of samples, is unknown and likely highly variable between systems. The correlation level set here in the *covariance matrix simulations* was between −0.88 to 0.89 (Appendix N). The mean empirical correlation of the samples in the *ecological simulations* for set-parameters simulations when there is a positive interaction is 0.168 and the mean for negative interactions is −0.170 for simulations with 10 species. For 100 species, it is 0.118 for positive and −0.105 for negative interactions.

We also explored several additional *alternative covariance matrix simulations*, which have a different simulation mechanism specified in Appendix G; these include some simulation sets with a lower and intermediate level of correlation. In one case, we ran an additional test of linear and logistic regression on *alternative covariance matrix simulations* where 100 species were associated in 50 independent pairs and the correlation level was set at 0.5. In these simulations, 250 samples were sufficient for linear regression without covariates to classify the samples nearly perfectly according to ROC, with logistic regression as a close second (Appendix G). Other methods were not tested on these data, although based on performance with large sample sizes in the *covariance matrix simulation* model, we believe that they would perform similarly well under this idealized scenario. Many factors, including covariate responses, were not modeled here so we believe this is a less realistic scenario. We also ran *alternative covariance matrix simulations* with correlation values set at 0.1 to correspond more closely to the correlation of the *ecological simulations*. In these cases, 100 samples was not sufficient for any method to perform well, but they performed well with 10,000 samples (Appendix G).

### Information about species abundance has a mixed effect on inference results

Some methods tested here take sedaDNA read abundance as input, which is considered a proxy for relative species abundance or relative biomass. Others take presence-absence data, which is derived from read abundances by setting a threshold for how many reads indicates species presence. Using presence-absence data is justified if read abundances are biased and therefore are not a good proxy for organism abundance. Additionally, this can be considered a way to give low-abundance taxa a higher weight in the model [43]. We consider these methods to fundamentally take the same data, but process it differently.

We find that information about species abundance improves inference of species associations in the *ecological simulations* with set parameters, but not on average across all sets of parameters when they are set at random. In the simplest case, we compare logistic regression (presence-absence data) to linear regression (read abundance data). For the simulations where any methods perform better than a random classifier, linear regression performs better than logistic regression according to ROC curves in the set-parameter *ecological simulations* (Figure 5, 6, 7, 8). EcoCopula is the only other method that flexibly takes presence-absence or read abundance data. The performance of EcoCopula varies, but does not seem to change based on data type for the *ecological simulations* (Figure 4).

On the other hand, for the *covariance matrix simulations*, in several cases we see that the performance of logistic regression (presence-absence data) exceeds that of linear regression (read abundance data) (Figure 4, 5, 7). Additionally, EcoCopula in several cases has a lower FDR with presence-absence data than with read abundance data (Figure 4). In the *covariance matrix simulations*, read abundance data is not a direct proxy for species abundance since no species abundances were simulated. More assumptions are violated by the count data than the binary data in this case, which may explain differences in performance.

Here we have not added species-level biases to simulated read abundance, which means that read abundance is correlated to species abundance in the *ecological simulations*, although often with significant levels of noise. More biases may exist in read abundance in real data, so further investigation is needed to determine the circumstances under which sedaDNA read abundances are a good proxy for species abundances.

### Measuring environmental covariates can improve inference performance, but it can also have a negative effect when there is multicollinearity

In the *covariance matrix simulations* where there is a covariate effect, we see that the only cases where the FDR is low is for the methods that correct for the effect of covariates. That is because without correcting for the covariates, the methods are detecting shared responses to similar environments. This could still be defined as an association in some studies, so this is mainly a comment about interpretation (Figure 3). For example SpiecEasi and SparCC cannot separate the environmental covariate effects from residual correlation between species because they do not accept covariates as input. This demonstrates that, as expected, these methods are often detecting shared responses to the environment rather than species interactions. We see the same effect for other methods when the covariates are unobserved (Figure 4). However, even in the cases where all covariates are observed perfectly, the false discovery rate of interactions is still much higher than the cases with no covariate effects in the simulation (Figure 4). Even if covariates are measured and included as input in the inference method, they can still cause errors. For example, in the *covariance matrix simulations* with covariate effects, the methods that take the Poisson distributed read count data still perform poorly even when they attempt to correct for the environment, even though all variables are measured. We believe this is because of violations of assumptions of the methods. In support of this conclusion, linear regression performs relatively well on the intermediate latent Gaussian variable *z*_*j*_(***s***, *t*), but poorly on the Poisson reads derived from that variable (Appendix P). Another factor is that drawing the Poisson variable adds higher variance and thus the signal to noise ratio is lower, but since this issue persists at large sample sizes (if anything, it is exacerbated), we believe it is more likely attributable to mismatching assumptions.

On the other hand, contrary to expectations, in the *ecological simulations* inference of species associations is often worse with environmental covariates included in the model than without them (Figure 5, 6, 7, 8). We would generally expect the inclusion of many covariates to cause loss of statistical power (this would not affect ROC curves), but improve accuracy by correcting for confounding variables [42]. However, if many covariates are included in a model and they do not affect multiple species, correlations caused by random fluctuations in the data can cause false discovery rates to be higher (Appendix H).

Additionally, in the set of simulations with set parameters, there is a high level of correlation between some of the environment covariates and the species that they affect. In the methods that take both species read abundances or occurrences and covariates as inputs, this can cause issues with identifiability between the effect of the covariate and the effect of the species (Appendix I). Therefore, it is important to check for collinearity between species and covariates if both covariates and species are going to be included as explanatory variables in these models.

### SPIEC-EASI is highly sensitive to assumptions about number of species

SPIEC-EASI assumes that the number of species is large, although the actual number of species required is not specified. There is an approximation in their model that is more accurate as the number of species gets large [8]. We find that at the 5-10 species level, SPIEC-EASI is highly sensitive to this assumption according to ROC curves when looking at direct associations (Figure 5). With 50-100 species, this sensitivity disappears, and it diminishes when looking at indirect associations (Figure 6, 7). Specifically, with below 10 species, we find that SPIEC-EASI seems to have strong evidence for actual negative associations between species and strong evidence against actual positive associations between species (Appendix J). This effect is not observed after model selection because the model never selects the regularization parameter in this region of the ROC curve (Figure 5, 6). Therefore, the false discovery rates after model selection are not noticeably affected (Figure 4).

### Regression performs similarly to other methods with the same data

Contrary to expectations, we find that in most scenarios, linear and logistic regression perform, on average, similarly to other methods with the same data input (Figure 5, 6, 7, 8). We do not interpret this to mean that regression is the correct model for this data, but rather that the more complex methods do not improve modeling of the data structure in these scenarios. Each of the more complex methods attempts to account for different aspects of the data better than linear or logistic regression, but we find that for a variety of simulation scenarios, they do not succeed in better modeling the data generating process. It may be the case that under certain circumstances, each model does perform better, but we find that on average across many scenarios, no method recovers associations consistently better than regression. Additionally, regression methods are much faster than any of the other methods. In the case of the *covariance matrix simulations*, very few of the assumptions of logistic regression are violated. Therefore, it is expected that it would perform well given enough data. In the simulations shown here, we use the Benjamini-Hochberg method for false discovery rate correction. However, we also explored the use of no correction (less conservative) or the Bonferroni correction (more conservative). We found that the overall trends were similar, although the exact values of the false discovery rates vary considerably (Appendix O). The ROC curves would not change based on the correction method. Rather, each correction method chooses one point on the corresponding curve. Linear and logistic regression will not work with the number of predictors (number of species plus covariates in this case) being greater than or equal to the number of samples. Therefore, for simulations with 100 species, the 100 samples case was omitted for the regression methods. Additionally, for logistic regression, when the number of predictors is close to the number of samples, we encounter numerical problems because the samples provided can be separated perfectly by the regressors. Therefore, results for logistic regression with 100 species and 250 samples are also not shown.

For methods that take presence-absence data, SDM-INLA performs best when averaged across random simulation parameters at a high number of samples in the *ecological simulation*, and performs similarly to other methods in the *covariance matrix simulation* (Figure 5, 7). However, this result does not generalize to our set-parameters *ecological simulation*, where logistic regression performs better (Figure 5, 7). The main difference between these models is that SDM-INLA models spatiotemporal autocorrelation. The effect of this may depend whether a dataset has spatiotemporal structure and whether this structure is correctly modeled by the SDM-INLA method.

In simulations with random parameters, we see less separation of the different methods in the ROC curves than when the parameters were set at a single level (Figure 5, 6). This is the effect of averaging performance across many different ecological scenarios. In some ecological contexts, some methods may perform better than others, but this may not be consistent across all contexts. As expected, the variance of the results within a method when parameters were varied was much higher than when parameters in the simulation were held constant.

### Many dimensions of simulation parameter space affect the success of regression models in predicting direct, symmetric interactions

In the *ecological simulations* produced with random parameters, there was a great variety of resulting false discovery rates between different parameter sets. In order to better understand how the simulation parameters are affecting the success of the inference models, we created an additional 1000 data sets with random parameters (drawn from distributions specified in Table 4), and used these simulation parameters as predictors for a random forest model which predicts false discovery rates of direct, symmetric interactions. Due to the large number of simulations in this set, this was only performed for logistic and linear regression, which are much faster than other methods. We believe this is justified since no other methods consistently outperformed these methods in tests on smaller simulation sets, though we do not know if the same simulation parameters would affect the performance of all methods. Random forest models were verified by testing for predictive accuracy on an independent test set of 100 simulations. RMSE for the test sets were evaluated against a naive predictor of FDR (median of observed FDRs). The random forest predicts FDR between 8% and 15% better than the naive predictor (Appendix K), indicating that the simulation parameters have some predictive power, though a lot of the variation in FDR is not well predicted.

For 10,000 samples, three simulation parameters seem to affect FDR significantly for logistic and linear regression. For larger *r* (population growth rate for all species) models tend to have lower FDR (perform better) (Appendix L). Larger population growth rates will cause the populations of species to change more rapidly and to reach their carrying capacity faster, causing less lag between a change in environment and the change in the species population. Non-equilibrium dynamics caused by lag between changing conditions and population growth can cause temporal and spatial autocorrelation not accounted for by covariates. This violates the assumption of conditional independence of the samples in linear and logistic regression and can lead to a loss of information.

With a larger migration radius (*d*), the models tend to have a higher FDR (perform worse) (Appendix L). In this simulation model, migration rates are not affected by species interactions and may obscure the effect of one species on another.

The third simulation parameter that seems to affect false discovery rate in this set of simulations is the individual sampling probability (*p*_*n*_) (Figure 9). When this parameter is higher, FDR tends to be higher (Appendix L). This effect is stronger for logistic regression than linear regression. This is a counter-intuitive result because we would expect a higher probability of sampling individuals to result in better performance of the models. However, we also observed that high observed rate of species presence can result in difficulty with inference because there is less power to infer interactions if species are present almost everywhere, which we believe to be the cause of this effect (Appendix M).

**Figure 9:**
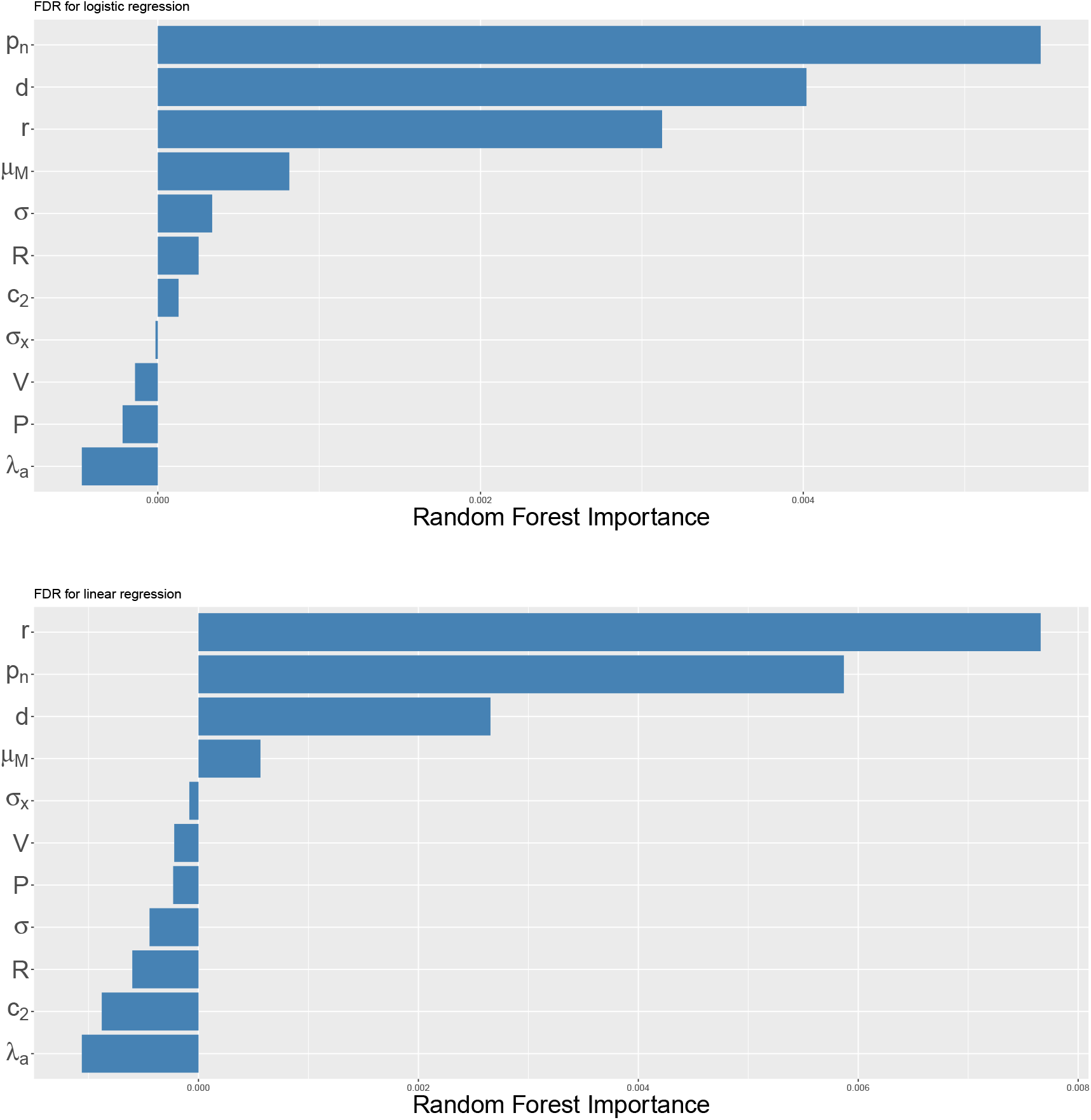
Importance of predictor variables in random forest predicting FDR as a function of simulation parameters. Analysis performed with linear and logistic regression of each species’ data as a function of all other species data. Simulations: *ecological simulations* with 10 species, random parameter settings, and 10,000 samples per simulation. Benjamini–Hochberg correction was applied with false discovery control level of 0.05, and FDR was evaluated for direct, symmetric interactions. Random forest was trained on results from 1,000 simulations.

## Discussion

We have tested a range of methods which have been used to detect associations between taxa from sedaDNA data. We simulated data under a variety of models, including a simple model where all data points are independent over time and space, and a custom ecological model that leverages principles of ecological theory to create more realistic data. Explicitly simulating population dynamics through time in the presence of a changing abiotic environment will create patterns in the data that mimic those in nature. We find that in all but the most idealized scenarios, false discovery rates of species associations are high for all methods tested.

Learning how past ecosystems have changed in response to environmental change will be an essential part of understanding and mitigating present-day global climate change and making informed resource-management decisions [25]. Biotic interactions are a key piece of this puzzle [4] and sedaDNA is a new frontier in understanding ecological interactions. It has the potential to uncover interactions within and between trophic levels, model species distributions over large spatiotemporal scales, and show how biodiversity has changed over periods of massive environmental change [1–3, 12]. Robust computational methods are needed to come to accurate and reproducible conclusions, and it is important to understand the relative performance of methods under different conditions.

Many species distribution models have been developed in ecology [11], but since these methods were developed with traditional ecological survey data in mind, we find that they are often not scalable to the number of taxa that is common in sedaDNA datasets. Many of these methods account for temporal and/or spatial information in the data, and allow for information about abiotic covariates [11, 12]. However, they do not account for the compositional structure of sedaDNA read abundance data. On the other hand, several methods have been developed specifically for sedaDNA data, and therefore account for non-independence due to compositional sedaDNA read data [8, 21]. However, many of the most commonly used methods were developed specifically for microbiome sedaDNA data and therefore assume large numbers of taxa (or OTUs/ASVs) in a single data set. We find that SPIEC-EASI is very sensitive to this assumption, exhibiting very poor performance on data with small numbers of taxa (10 or fewer). This is important, even in the sedaDNA field, because depending on the taxa under study and the taxonomic level of assignment, many studies may want to do similar analyses with small numbers of taxa [12, 14].

It has been previously established that inferring species interactions from spatial presence-absence data is a challenging statistical problem, and that there are many potential causes for false inferences from these data [42]. It has also been documented, theoretically for co-occurrence (presence-absence) data [42] and empirically for read abundance data [8], that high sample sizes are needed to accurately estimate associations between taxa. The challenge is in part because a large number of taxa create an even larger set of potential interactions [8], but also due to the complexity of the ecological networks involved and the abiotic factors that influence them, among other reasons [42]. From a mathematical perspective, for linear regression with all assumptions met, when the correlation is 0.1 (close to the average in our *ecological simulations*) using a p-value cutoff of 0.05 with Bonferroni correction, the necessary sample size to detect 80% of true interactions is nearly 2000 samples for 10 species and nearly 3000 samples for 100 species (Appendix F). If the correlation induced by species interactions is higher, the expected sample size needed would go down (Appendix F), which we also observe for simulated data (Appendix G), so lower samples sizes may be sufficient under some circumstances. However, even in these cases, false discovery rates may remain high due to model misspecification. Since very few studies using sedaDNA currently have more than a few hundred sampled points, high sample sizes are currently unrealistic, but we are hopeful that future data will rise to meet this challenge. Additionally, shared responses to the environment may cause associations which are separate from direct interactions between species [9], and therefore in real data we would caution users to interpret inferred associations carefully. We also observe in this study that even when the environmental variables are fully observed and statistically independent across time and space (thus minimally violating the assumptions of the methods), they still invariably cause higher false discovery rates than were seen in the *covariance matrix simulations* without a species response to the environment.

False discovery rates reported here depend heavily on the calibration of various methods (such as model selection or choosing a p-value cutoff). FDR reported here reflects one point on the corresponding ROC curve, which measures the performance of the model independent of calibration. Since calibration is an intrinsic part of these methods, we consider these calibrated values to have substantial significance. The dramatic differences in calibrated FDR values between different versions of the same method and between methods is often more a consequence of the calibration than the underlying model. For example, the ROC curves for all methods may look similar across all models in many cases, reflecting similar success of the underlying model, but the average FDR varies considerably due to poor calibration of some methods compared to others. Even within the same method and simulation set, the calibration can cause very different results depending whether covariates are included. Additionally, the number of total inferred interactions varies dramatically between methods and simulations, so some average FDR values may have high variance between simulation runs (Appendix B).

Sample size is not the only potential concern, since we observe high false discovery rates in our *ecological simulations* and in the *covariance matrix simulations* with covariate responses even with high sample sizes. With very high sample sizes, statistical methods should perform well when the data meets their assumptions perfectly, but when the true data generating process is different from the assumptions of the method, the error rate may not go down with more data. The assumptions about the data generating process vary across methods. For example, none of the methods tested here account for uncertainty in covariates. Some are parametric methods that rely, for example, on residuals being independent and normal [16]. Many of them do not account for residual autocorrelation in space and/or time, which may be caused either by non-equilibrium distributions or by autocorrelation of an unmeasured abiotic factor [16, 44]. Some methods cannot detect non-linear interactions between species and covariates or between species. These are all examples of assumptions that we have violated to varying degrees in the simulations. We have shown here that given enough samples, some methods perform moderately well with some assumptions violated, but the only case in which any method is able to reliably recover interactions with a low false discovery rate is under an extremely idealized scenario.

Many simulation studies use data that assumes the same data generating process as the methods they are testing. Since real data is not expected to follow the assumptions of the methods exactly, this procedure should be expected to overestimate performance. As in all simulation models, we make many assumptions about the data generating process, but in our case, some of the simulations significantly deviate from the assumptions of the inference methods. In our *ecological simulations* we assume that populations in this simulation will grow according to a logistic growth curve with a changing carrying capacity through time. This may not be the optimal way to calculate growth rates for some environments and species [45]. Additionally, since actual ecological network structure is likely variable in different systems [6], we assign interactions between species randomly in our *ecological simulations*. This is likely a conservative assumption because graphs with evenly distributed neighborhoods are easier to recover than those with large hub nodes [8]. In the *covariance matrix simulations*, interactions are instead organized in clusters of interacting species. Differences in performance due to structure of the interaction networks between species have been explored [25], and the inference methods discussed here are often used to estimate overall characteristics of the network [8], so these assumptions may be significant. Additional assumptions of our *ecological simulation* model include assuming that some species traits and data characteristics are constant across time, space, and species, which is certainly not always the case. For example, we expect that covariate uncertainty, average read abundances, and species detection rates may vary in space and time and across species [46, 47]. Our simulation model also assumes that interactions between species are constant over time and space, which may not be true over evolutionary timescales. All of these assumptions are conservative, but we recognize that we also made assumptions about the data that are less conservative. For example, strong species interactions in real data may induce higher levels of correlation than we see in our models. Additionally, other characteristics of the simulation may create unrealistic dynamics that affect the performance of the inference models. However, within our chosen simulation framework, we attempt to simulate many ecological dynamics by varying the parameters widely in our simulations. Additionally, we test the methods under a simpler simulation framework (*covariance matrix simulations*) that is not based on ecological theory but is expected to minimally violate assumptions of the inference methods.

The input to some models was sedaDNA read abundances, whereas other methods take only presence-absence data. EcoCopula was the only model that takes flexible input. We find that information about species abundances improves performance of models at detecting species interactions in our *ecological simulations*. On the other hand, in the *covariance matrix simulations*, we see more mixed results. In a few cases, methods with presence-absence data perform better than their read abundance counterpart. We believe this could be due to differences in the distribution assumed by the models and the actual distribution of read abundances. For example, linear regression assumes the residuals are Gaussian and the actual read abundances are Poisson distributed. Additionally, the Poisson abundances have a second source of random variability from drawing the latent Gaussian variable first and then the Poisson variable for the reads. We believe that this may be realistic, since the read counts may also introduce extra variability in the data as compared to treating the data as presence-absence data. Additionally, in real data, read abundance data may be biased by taxon, which may introduce confounding factors that result in spurious correlations. For example, many factors may create biases such as species-specific DNA deposition rates [29], PCR bias [48], or bias in species assignments [47]. In our simulations, we did not include per-species bias in rates of DNA deposition, so it is not surprising that simulated read abundances have more information about species interactions than presence-absence data, though we do not know whether this is the case in real data. These concerns would still be applicable to some extent in presence-absence data, as detection rates may also be biased across taxa, space, and time, but the influence of these biases may be lower. We also note that in microbiome studies, reads are not assigned to taxa but rather grouped into OTUs or ASVs, which may have different properties with respect to read abundance biases [24]. Further investigation will be needed to fully understand these effects and the circumstances under which it is effective to use read abundances as a proxy for species abundance.

The increasing availability of whole-community ecology data has enormous potential to uncover ecological dynamics, parameterize models, and make predictions about the future of ecosystems. However, data of this complexity must also be approached with caution, and benchmarking computational methods under a variety of scenarios is an essential step in understanding and interpreting results. Further investigations into the relative success of these methods and others will be needed for a complete understanding of their performance. For example, it is possible that these methods would be more successful at recovering overall network properties as opposed to specific interactions [8]. Additionally, many other methods exist that were not tested here due to differences in data requirements and outputs, including some mechanistic models, which may perform better under certain conditions [49, 50]. An area where sedaDNA has a lot of potential is in its ability to recover species distribution data across a wide range of taxa, including microbes, plants, and megafauna, from the same samples. However, tools to analyze data on differing taxonomic scales have been developed with differing assumptions (perhaps for good reason). In order to fully take advantage of the potential of these data, we will need analysis frameworks that are able to handle many different ecological scenarios.

## Appendix

**A JSDM-MCMC and SDM-INLA methods time to run**

**B Number of Inferred Interactions**

**C Convergence of JSDM-MCMC**

**D SpiecEasi number of inferred interactions versus number of samples**

**E Results using different methods of counting false positives**

**F Sample size versus correlation for linear regression**

**G *Alternative covariance matrix simulations***

**H Effect of adding many unnecessary covariates**

**I Multicollinearity**

**J SPIEC-EASI performance on low number of species**

**K Random forest performance on test set**

**L Random forest ICE plots**

**M Actual interactions cause higher percent presence on average in *ecological simulation*, but logistic regression infers interactions more for species with close to 50% presence**

**N Distribution of covariances for *Covariance Matrix Simulations***

**O Comparison of BH and Bonferroni correction**

**P Linear regression on Gaussian z-values**

## Notes

### Competing Interest Statement

The authors have declared no competing interest.

### Summary of Updates

Clarification added. Simple (covariance matrix) simulations updated to simulate count data in a different way.

