## Appendix for "Challenges in detecting ecological interactions using sedimentary ancient DNA data"

#### Contents


**A JSDM-MCMC and SDM-INLA methods time to run** **2**

**B Number of Inferred Interactions** **2**

**C Convergence of JSDM-MCMC** **2**

**D SpiecEasi number of inferred interactions versus number of samples** **6**

**E Results using different methods of counting false positives** **8**

**F Sample size versus correlation for linear regression** **12**

**G *Alternative covariance matrix simulations*** **14**

**H Effect of adding many unnecessary covariates** **18**

**I Multicollinearity** **19**

**J SPIEC-EASI performance on low number of species** **20**

**K Random forest performance on test set** **23**

**L Random forest ICE plots** **24**

**M Actual interactions cause higher percent presence on average in my simulation, but logistic regression infers interactions more for species with close to 50% presence** **27**

**N Distribution of covariances for *Covariance Matrix Simulations*** **29**

**O Comparison of BH and Bonferroni correction** **30**

**P Linear regression on Gaussian z-values** **31**

References **34**

### Appendices

#### A JSDM-MCMC and SDM-INLA methods time to run

INLA with 100 species: For one simulation run with 100 samples, it took 12.63 hours using a maximum of 10 threads. To run this on even one set of 100 simulations, this would take several months.

For one simulation run with 10,000 samples, it took 101.02 hours using a maximum of 10 threads. To run this on even one set of 100 simulations, this would take over a year.

JSDM-MCMC with 100 species: For one simulation run with 100 samples, we ran this method for 10 days on 5 threads and it did not finish.

#### B Number of Inferred Interactions

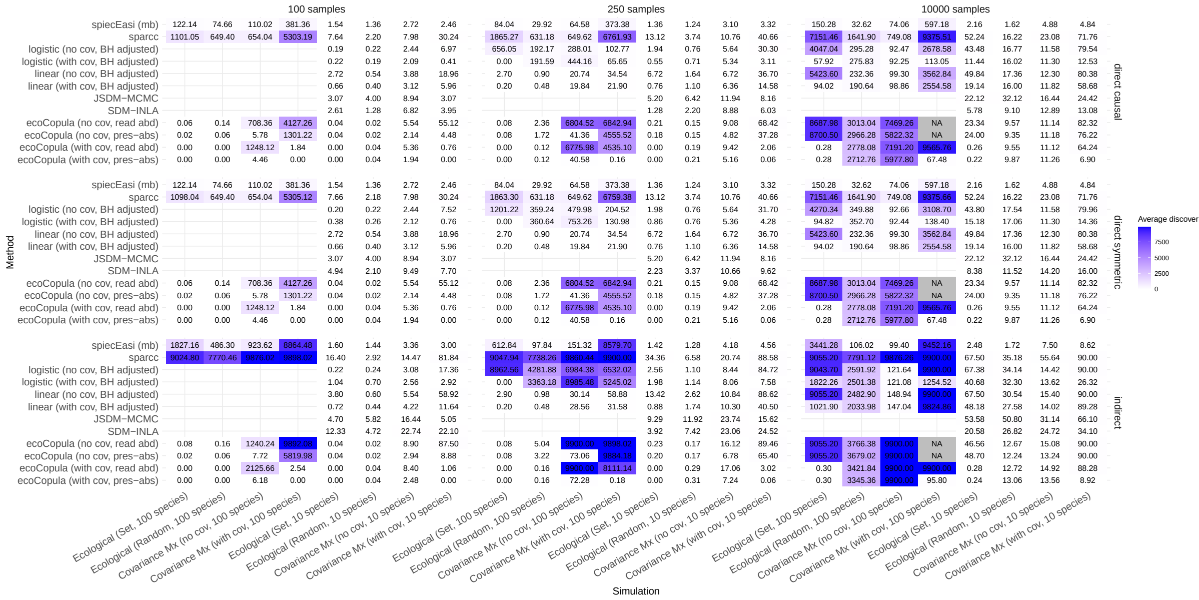


Figure S1: Discoveries per simulation on average across 100 simulations. When these numbers are very low, FDR estimates may have high variance.

#### C Convergence of JSDM-MCMC

The convergence rate of JSDM-MCMC is high so we ran 5000 iterations for each chain and discard the first 4000 iterations as burn-in. Here, we show trace plots for a representative example (interaction parameter between Species 2 and Species 3 for a randomly selected replicate of both the set-parameter and random-parameter ecological simulation with 10 species and 100 samples) and Gelman-Rubin statistics [1] for all parameters for a randomly selected replicate of set-parameter ecological simulation with 10 species and 100 samples.

| | 1 | 2 | 3 | 4 | 5 | 6 | 7 | 8 | 9 | 10 |
|---|---|---|---|---|---|---|---|---|---|---|
| 1 | | 1.49 | 1.01 | 1.38 | 1.67 | 1.34 | 1.08 | 1.36 | 1.56 | 1.46 |
| 2 | 1.49 | | 1.02 | 1.06 | 1.02 | 1.00 | 1.11 | 1.03 | 1.10 | 1.31 |
| 3 | 1.01 | 1.02 | | 1.20 | 1.13 | 1.48 | 1.19 | 1.35 | 1.12 | 1.16 |
| 4 | 1.38 | 1.06 | 1.20 | | 1.44 | 1.12 | 1.01 | 1.04 | 1.05 | 1.34 |
| 5 | 1.67 | 1.02 | 1.13 | 1.44 | | 1.04 | 1.31 | 1.11 | 1.88 | 1.00 |
| 6 | 1.34 | 1.00 | 1.48 | 1.12 | 1.04 | | 1.12 | 1.12 | 1.65 | 2.86 |
| 7 | 1.08 | 1.11 | 1.19 | 1.01 | 1.31 | 1.12 | | 1.03 | 2.10 | 1.03 |
| 8 | 1.36 | 1.03 | 1.35 | 1.04 | 1.11 | 1.12 | 1.03 | | 1.47 | 1.27 |
| 9 | 1.56 | 1.10 | 1.12 | 1.05 | 1.88 | 1.65 | 2.10 | 1.47 | | 1.31 |
| 10 | 1.46 | 1.31 | 1.16 | 1.34 | 1.00 | 2.86 | 1.03 | 1.27 | 1.31 | |

Figure S2: Gelman-Rubin statistics of correlations for a randomly selected replicate of set-parameter ecological simulation with 10 species and 100 samples

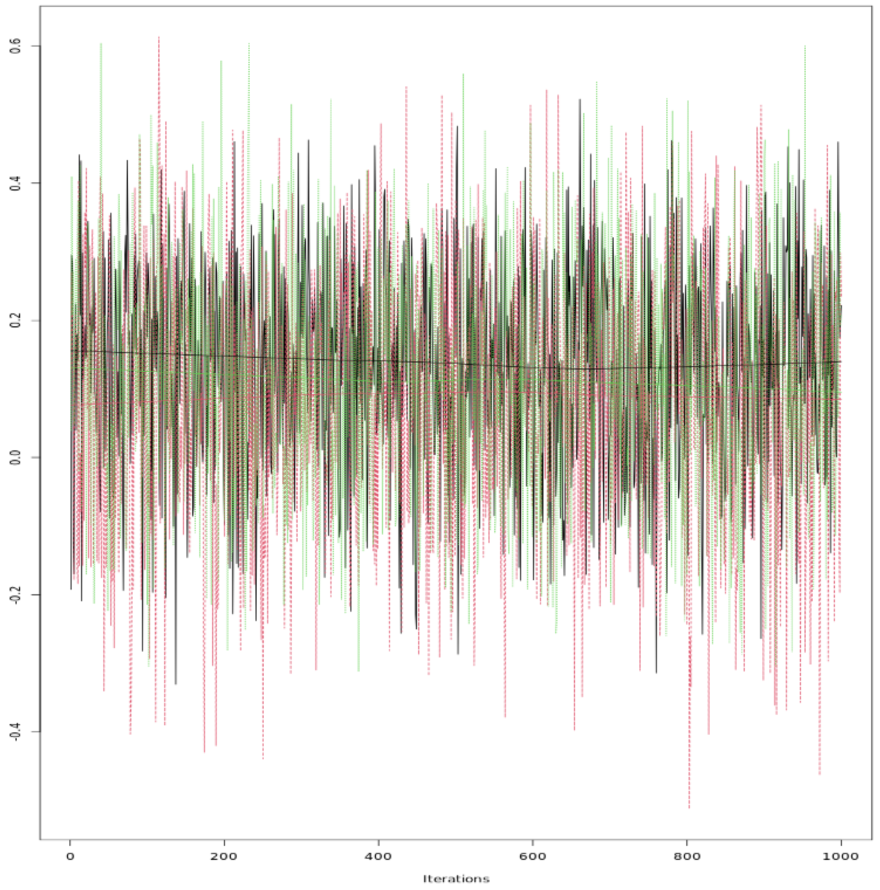


Figure S3: Trace plot of MCMC samples of the interaction parameter between Species 2 and Species 3 for a randomly selected replicate of the set-parameter ecological simulation with 10 species and 100 samples. Colors represent different chains, each with a different starting point.

| | 1 | 2 | 3 | 4 | 5 | 6 | 7 | 8 | 9 | 10 |
|---|---|---|---|---|---|---|---|---|---|---|
| 1 | | 1.04 | 1.00 | 1.14 | 1.15 | 1.06 | 1.39 | 1.07 | 1.01 | 1.29 |
| 2 | 1.04 | | 1.08 | 1.07 | 1.32 | 1.56 | 1.06 | 1.18 | 1.03 | 1.16 |
| 3 | 1.00 | 1.08 | | 1.33 | 1.02 | 1.53 | 1.00 | 1.08 | 1.05 | 1.08 |
| 4 | 1.14 | 1.07 | 1.33 | | 2.08 | 1.77 | 1.05 | 1.08 | 1.24 | 1.75 |
| 5 | 1.15 | 1.32 | 1.02 | 2.08 | | 1.29 | 1.45 | 1.22 | 1.31 | 1.02 |
| 6 | 1.06 | 1.56 | 1.53 | 1.77 | 1.29 | | 1.13 | 1.02 | 1.24 | 1.21 |
| 7 | 1.39 | 1.06 | 1.00 | 1.05 | 1.45 | 1.13 | | 1.27 | 1.39 | 1.01 |
| 8 | 1.07 | 1.18 | 1.08 | 1.08 | 1.22 | 1.02 | 1.27 | | 1.34 | 1.07 |
| 9 | 1.01 | 1.03 | 1.05 | 1.24 | 1.31 | 1.24 | 1.39 | 1.34 | | 1.92 |
| 10 | 1.29 | 1.16 | 1.08 | 1.75 | 1.02 | 1.21 | 1.01 | 1.07 | 1.92 | |

Figure S4: Gelman-test statistics of correlations for a randomly selected replicate of random-parameter ecological simulation with 10 species and 100 samples

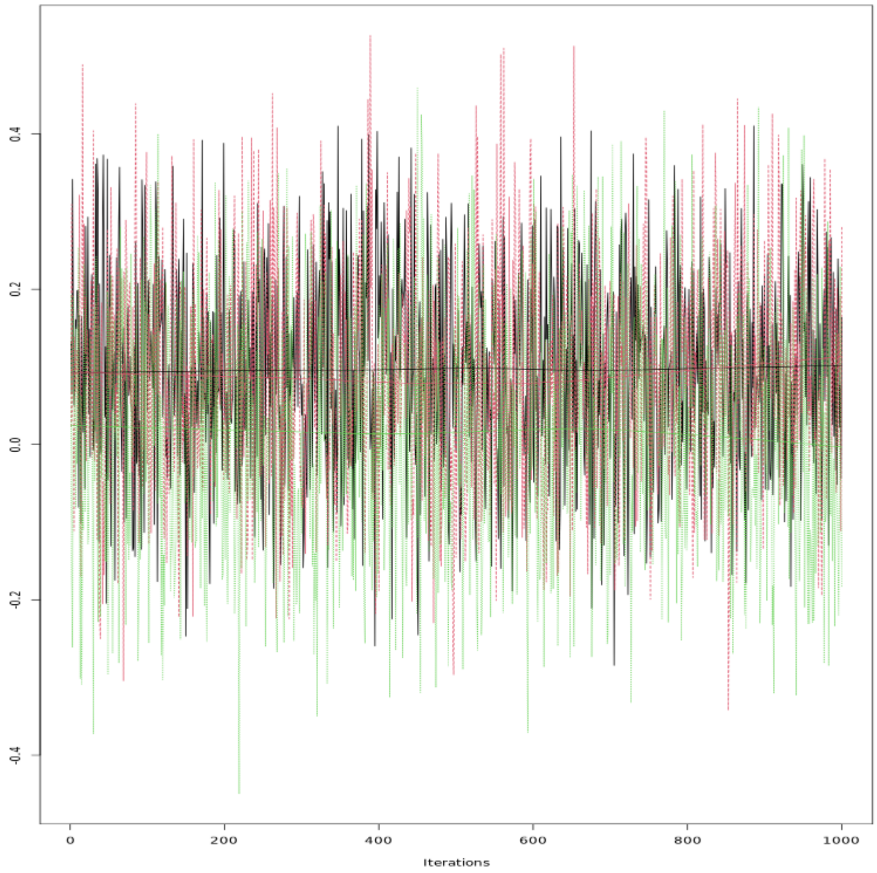


Figure S5: Trace plot of MCMC samples of the interaction parameter between Species 2 and Species 3 for a randomly selected replicate of the random-parameter ecological simulation with 10 species and 100 samples. Colors represent different chains, each with a different starting point.

### D SpiecEasi number of inferred interactions versus number of samples

Even with extremely low numbers of samples where there is unlikely to be enough information, SpiecEasi continues to infer many interactions using its default model-selection protocol. This is in contrast to many other methods that infer very few interactions at low sample sizes.

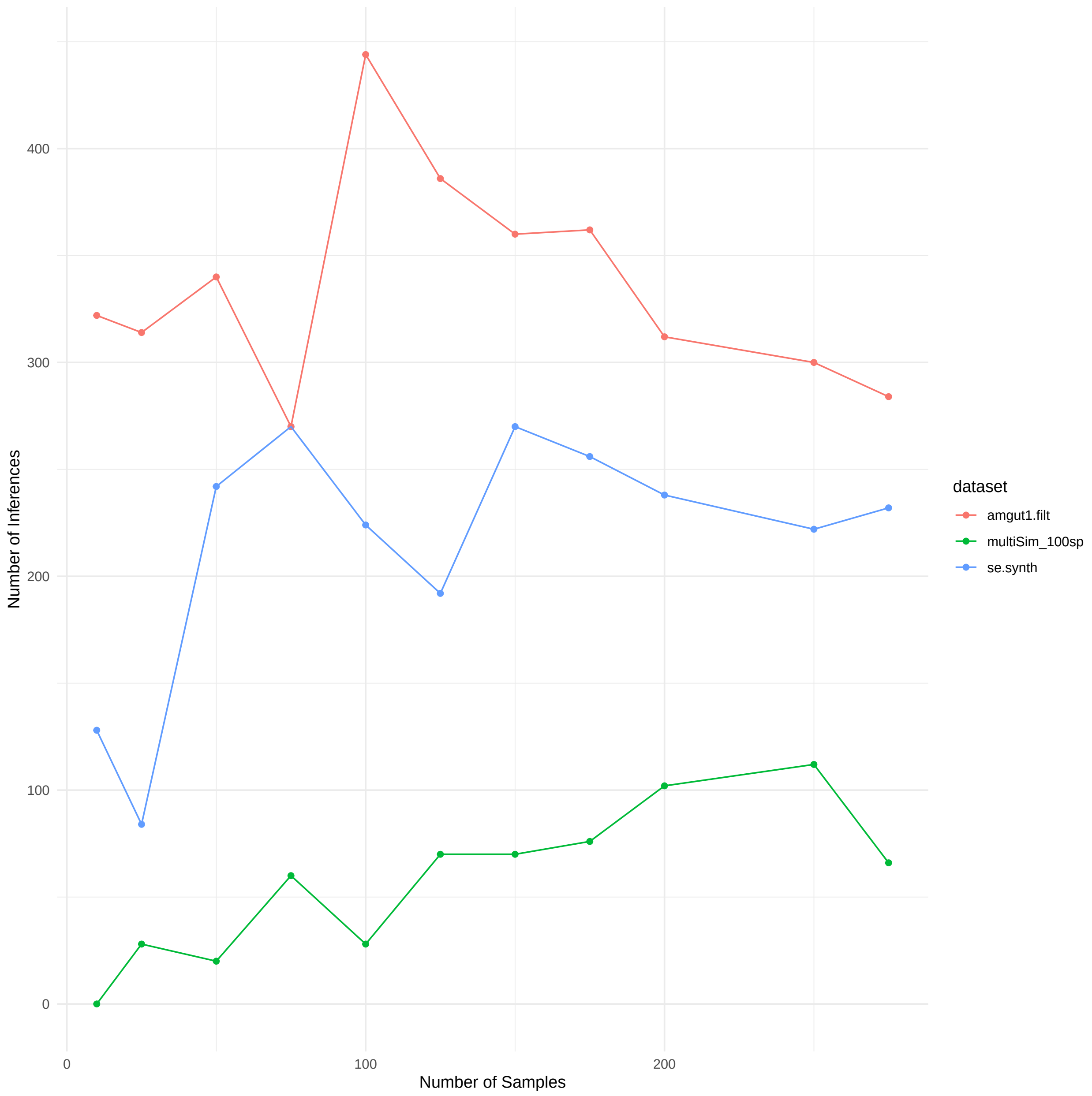


Figure S6: Number of inferences made using SpiecEasi on the example data included in their R package (amgut1.filt), the simulated data they provide as part of the SpiecEasi package (se.synth), and ecological simulation data with 100 species and set parameters (multiSim_100sp).

### E Results using different methods of counting false positives

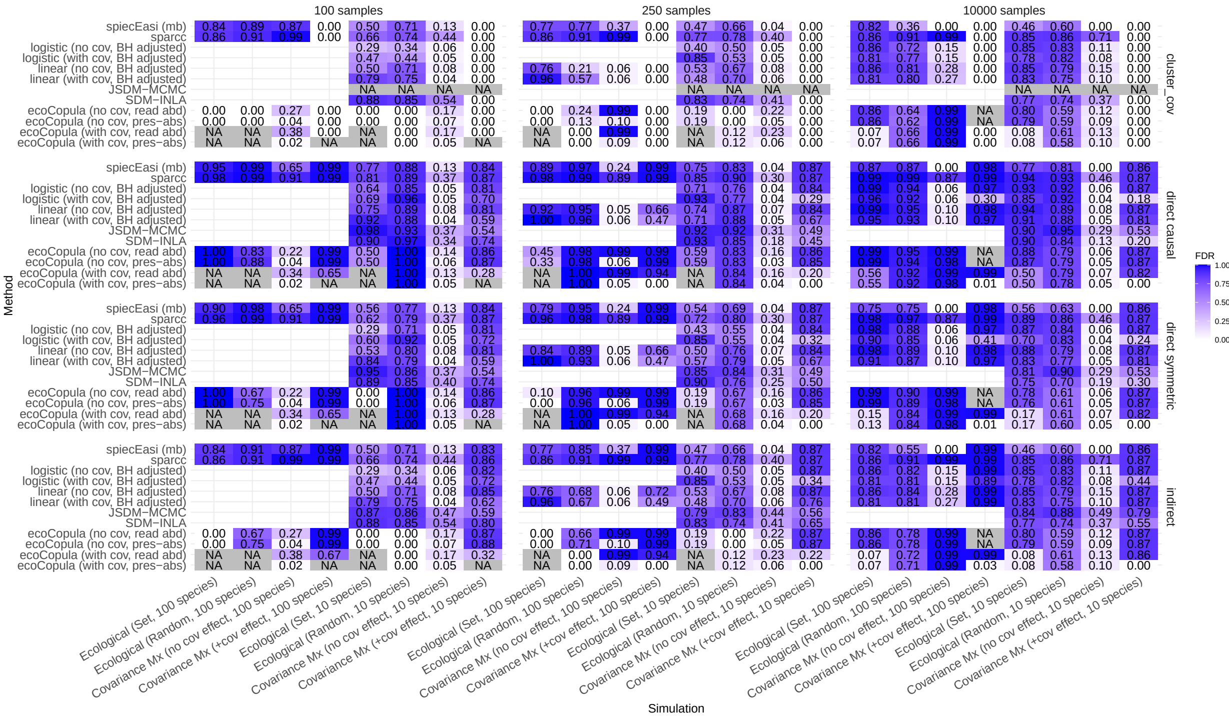


‘

Figure S7: Average false discovery rate (FDR) for all methods and simulation sets using all four methods of counting interactions. Cluster_cov is the method referred to in the text as indirect (covariate). This metric is not informative for the covariance matrix simulations because all species interact through covariates so the ground truth is that they are all connected.

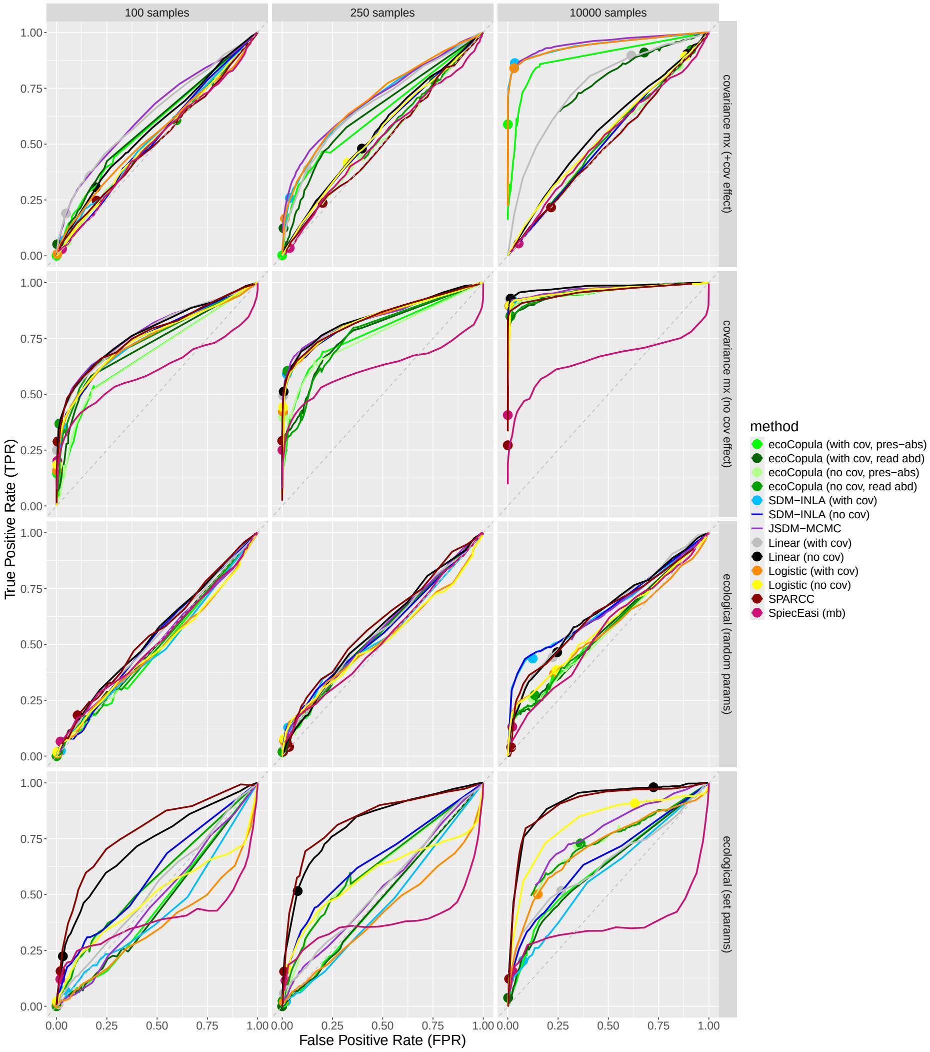


Figure S8: ROC curves for simulations with 10 species, measured using direct, causal interactions.

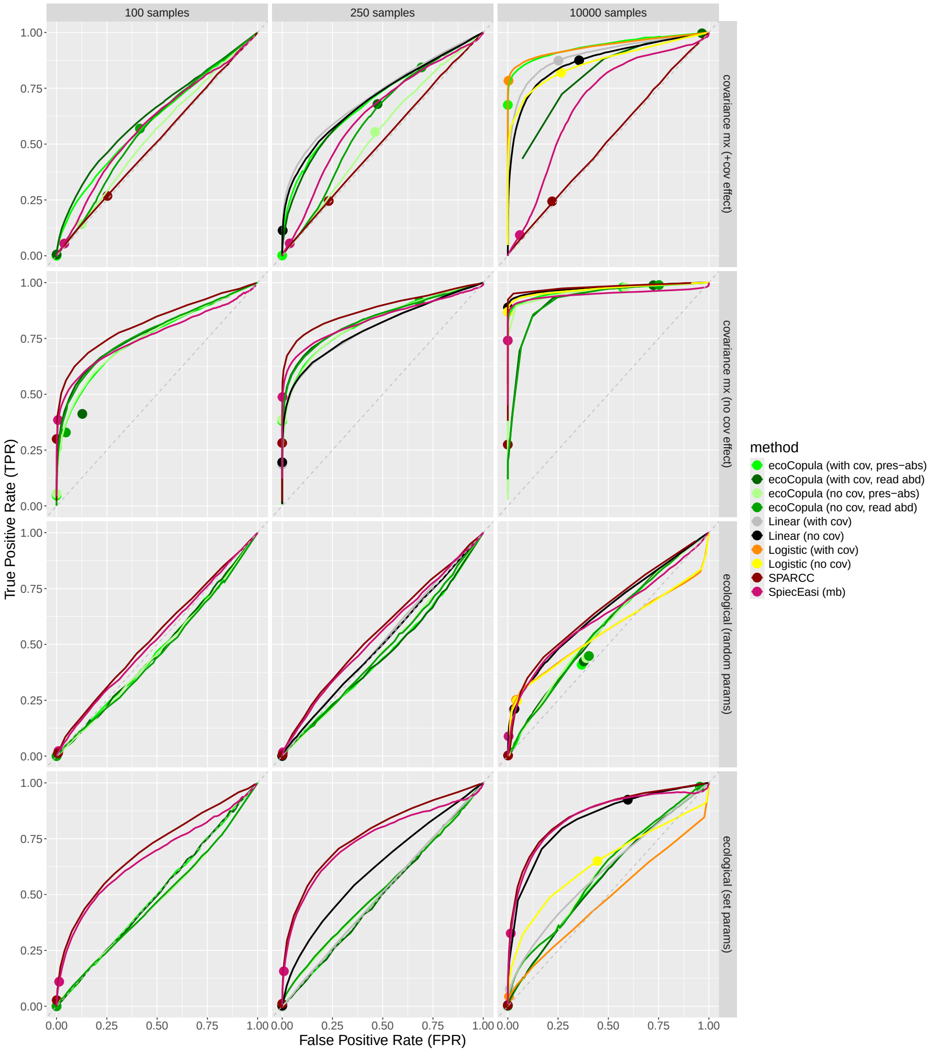


Figure S9: ROC curves for simulations with 100 species, measured using direct, causal interactions.

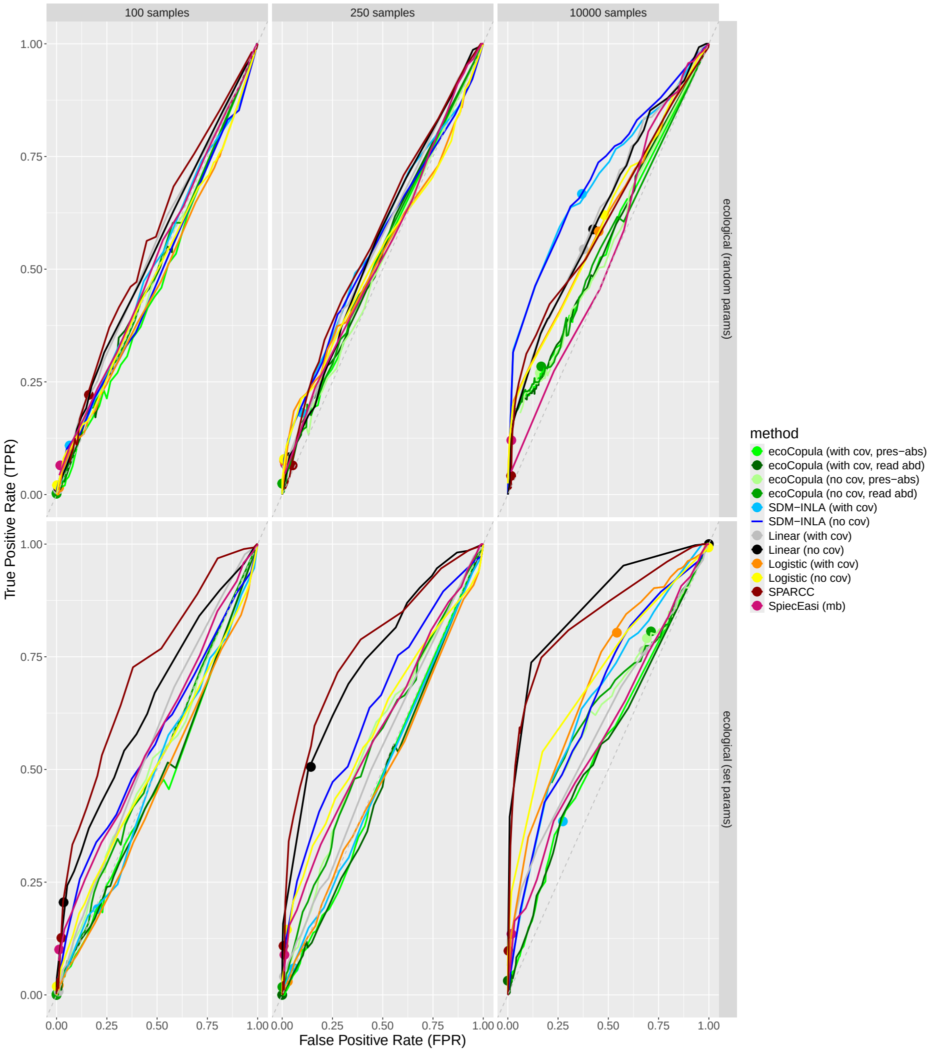


Figure S10: ROC curves for simulations with 10 species, measured using indirect (covariate-included) interactions.

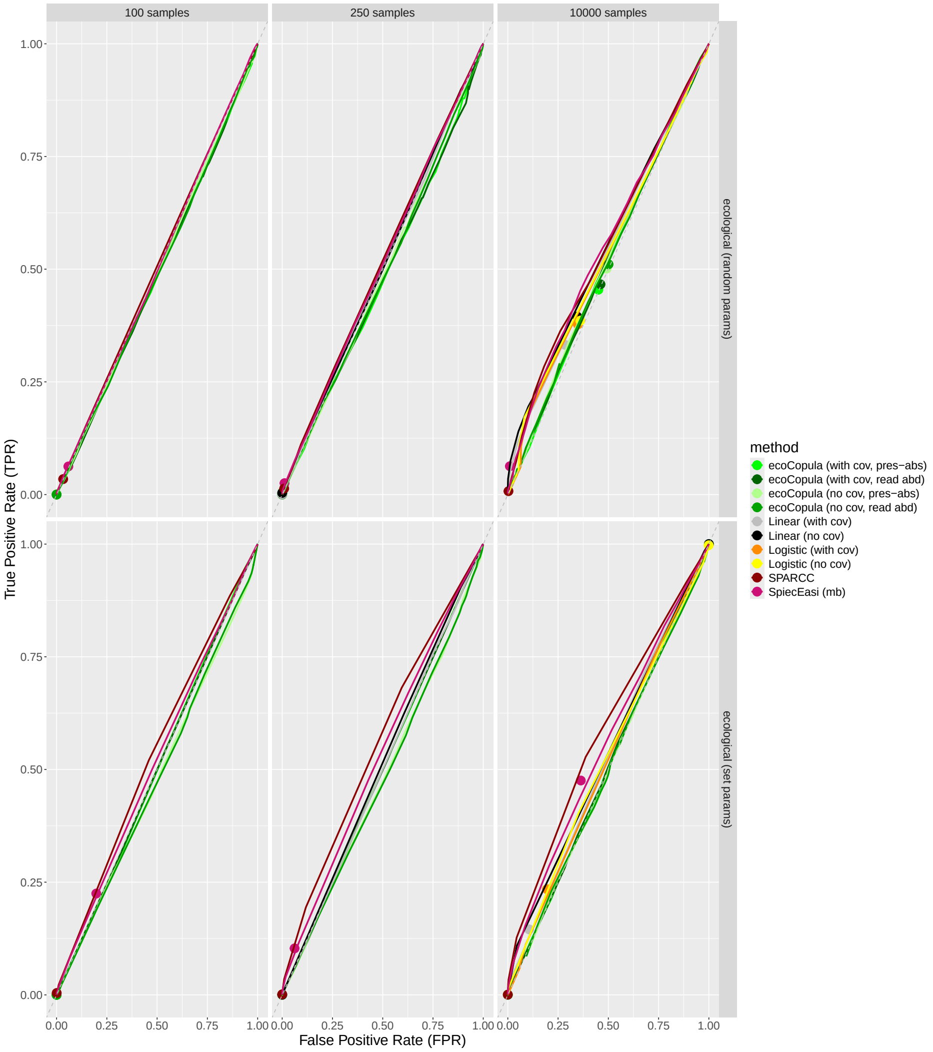


Figure S11: ROC curves for simulations with 100 species, measured using indirect (covariate-included) interactions.

### F Sample size versus correlation for linear regression

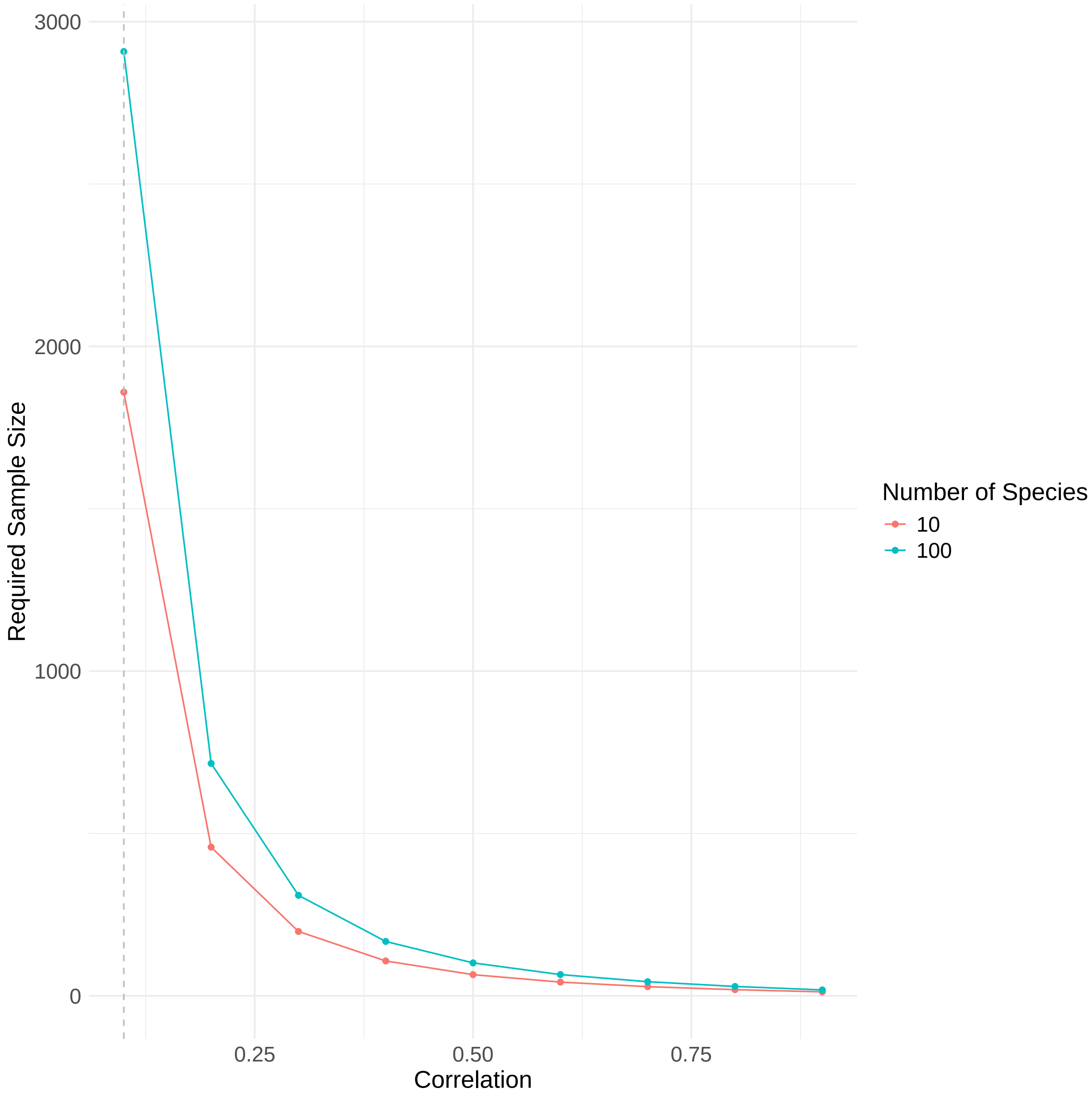


Figure S12: Necessary sample size to get power=0.8 for linear regression if all assumptions are met. This is for Bonferroni-corrected p-value cutoff of 0.05.

### G *Alternative covariance matrix simulations*

We also ran a different version of the covariance matrix simulations. The design was as follows:

Species read abundances are drawn from a mean-zero multivariate Gaussian truncated at 0. We use a fixed species interaction matrix, $\boldsymbol{\alpha}$, which is a sparse square matrix with dimension $J$ (number of species), and where the number in $\alpha_{j_1,j_2}$ is either 0, 1, or $-1$, indicating the sign of the interaction between species $j_1$ and species $j_2$. The covariance matrix for this set of simulations is defined in terms of $\boldsymbol{\alpha}$ in order to allow simulation of the same set of interactions with both the covariance matrix simulation model described in this section and the ecological simulation model described later. The correlation matrix for the Gaussian read abundances, $\Sigma_\alpha$, is defined as a $J$ by $J$ (same dimension as $\boldsymbol{\alpha}$) matrix with ones on the diagonal, 0.1 in all matrix entries where $\alpha_{j_1,j_2} > 0$ or $\alpha_{j_2,j_1} > 0$ (positive interaction), $-0.1$ in all matrix entries where $\alpha_{j_1,j_2} < 0$ or $\alpha_{j_2,j_1} < 0$ (negative interaction), and 0 elsewhere. This means that the expected correlation between the reads of species with positive interactions is 0.1 and the expected correlation between species with negative interactions is $-0.1$. The value of 0.1 is comparable to observed correlations between species that interact in the ecological simulation model, which is an emergent property of that model rather than a parameter (See Results). Read abundances are later multiplied by 100, which changes the variance and covariance but not the correlation.

Let $a(\mathbf{s},t)$ be the vector of simulated read abundances for all species at location $s$ and time $t$, then

$$\boldsymbol{a}^*(\mathbf{s},t) \sim MVNormal(\mathbf{0}, \Sigma_\alpha) \tag{1}$$

and

$$a_j(\mathbf{s},t) = \lfloor 100 \cdot \max(0, a_j^*(\mathbf{s},t)) \rceil \tag{2}$$

where $\lfloor \cdot \rceil$ denotes rounding to the nearest integer.

Due to the mean-zero truncated Gaussian, approximately half of the read counts are 0, which may be more or less realistic depending on the dataset. In this case, this is chosen to create optimal conditions for presence/absence data, as these methods lose power when percent presence is very low or high (Appendix M). Additionally, counts are multiplied by 100 to put them on the same order of magnitude as several example datasets, although we also recognize that this may vary between studies and depend on the level of classification of reads (e.g. OTU/ASV vs. species vs. family level) [2, 3]. Finally, data is rounded to the nearest whole number to create more realistic count data.

Presence/absence data, $y_j(\mathbf{s},t)$, is created by setting some threshold $R$ for the number of reads to determine species presence, as in Wang, et al., 2021 [4].

$$y_j(\mathbf{s},t) = \mathbb{I}\{a_j(\mathbf{s},t) > R\} \tag{3}$$

**Set 1 and 2: Interactions the same as ecological simulations, correlation 0.1, 10 and 100 species**

We simulated 100 datasets with 10 species and 100 with 100 species. In each simulation, 100 spatial locations arranged in a 10 by 10 grid were simulated for 10,000 time points, but the full simulated dataset was not analyzed. Although time and space are not included in this simulation algorithm, spatial and temporal labels were included in the final dataset to allow all methods to be tested. Each simulation was subsampled to 100 samples and 10,000 samples for analysis.

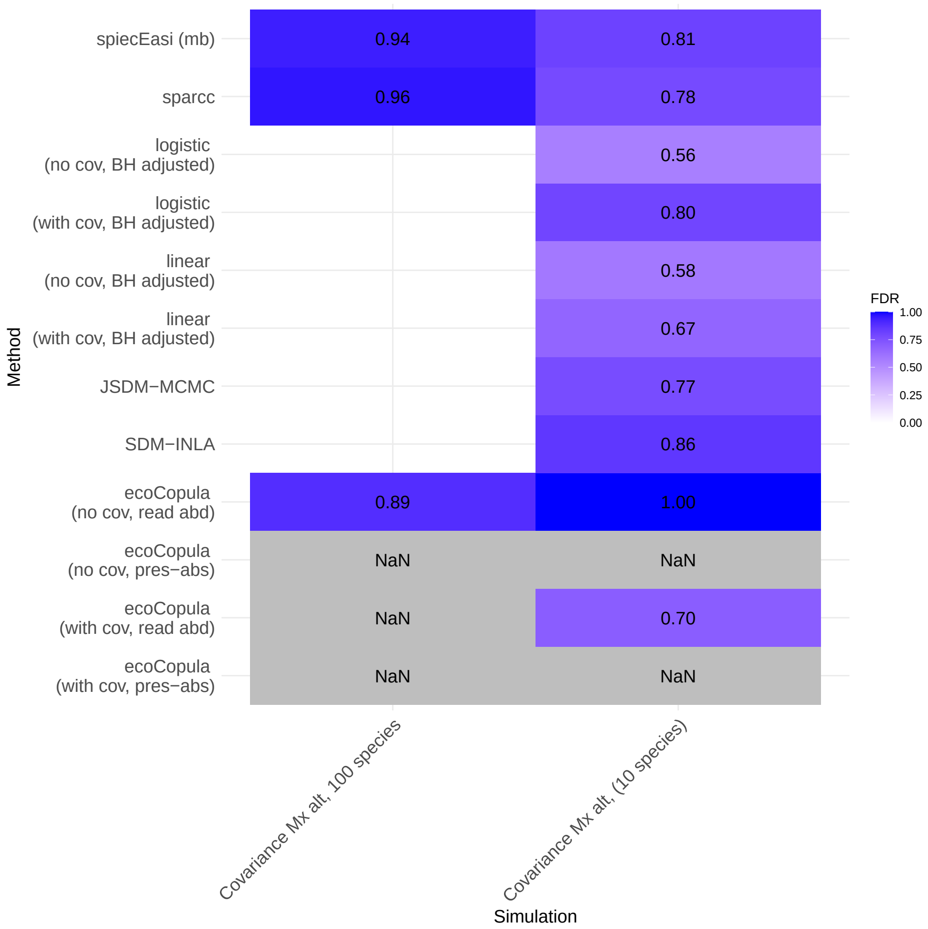


Figure S13: False discovery rates for alternate covariance matrix simulations (sets 1 and 2). 100 samples per simulation. FDR mode: direct symmetric, Benjamini-Hochberg correction.

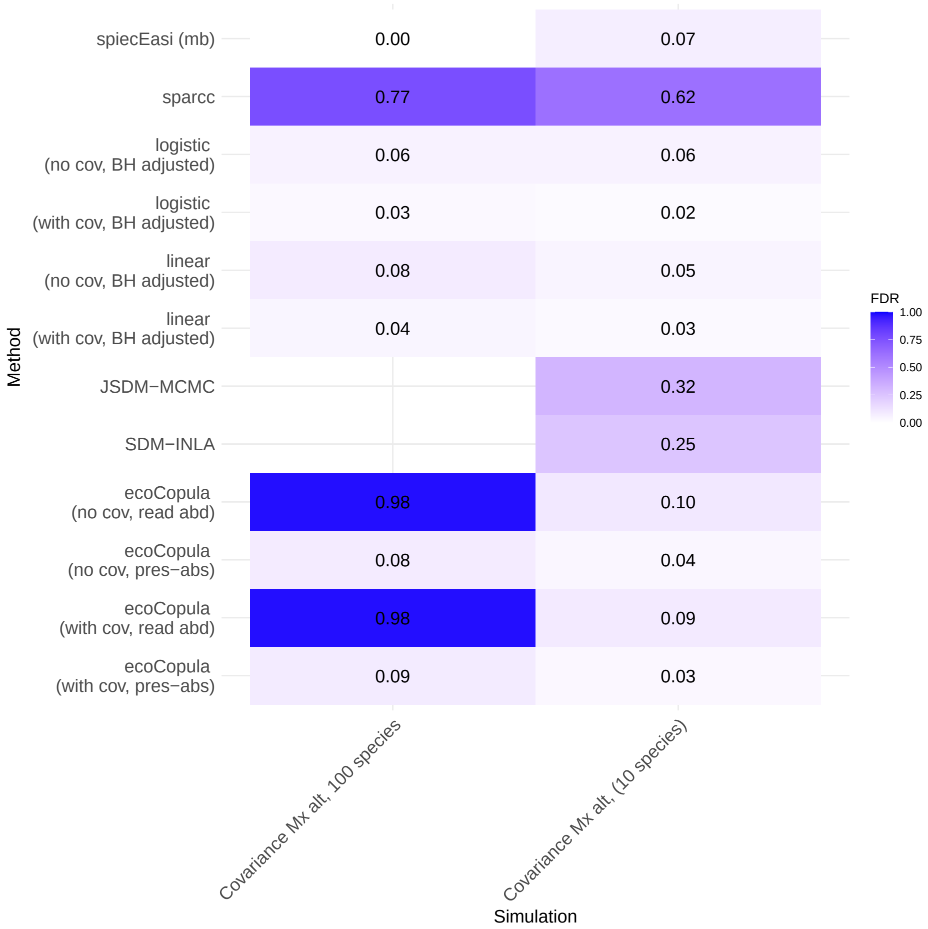


Figure S14: False discovery rates for alternate covariance matrix simulations (sets 1 and 2). 10000 samples per simulation. FDR mode: direct symmetric, Benjamini-Hochberg correction.

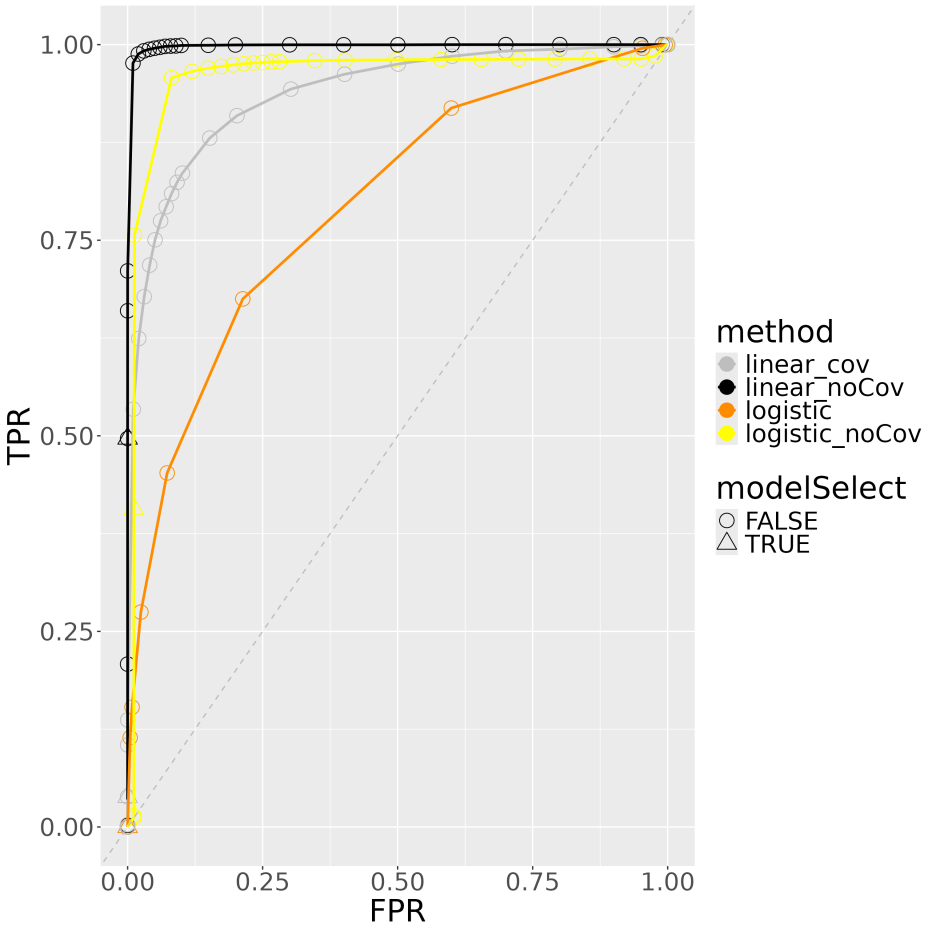


Figure S15: ROC curves for predictive success of species associations for linear and logistic regression using 250 samples. Simulation from alternate covariance matrix simulation (set 3). FDR mode: direct symmetric, Bonferroni correction.

**Set 3: Interactions in Pairs, correlation 0.5** Covariance matrix simulations used with correlation set at 0.5 for 50 pairs of interacting species. 100 species total. All covariances set to 0 except pairs that were set to 0.5.

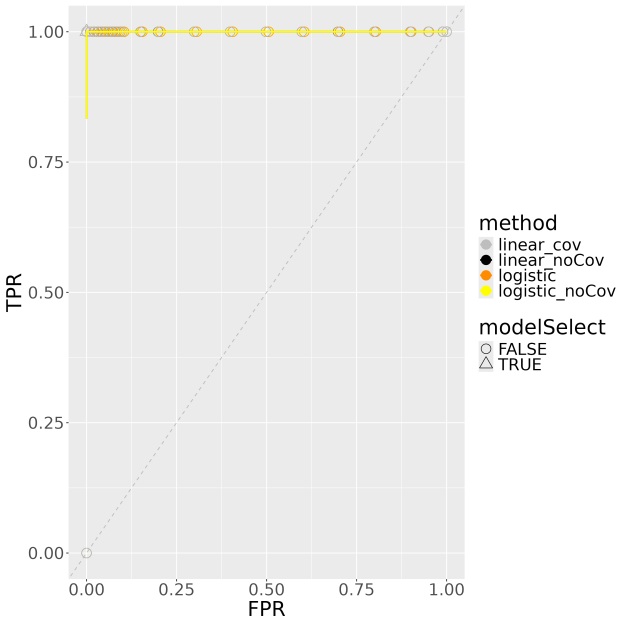


Figure S16: ROC curves for predictive success of species associations for linear and logistic regression using 10,000 samples. Simulation from alternate covariance matrix simulation (set 3). FDR mode: direct symmetric, Bonferroni correction.

### H Effect of adding many unnecessary covariates

Simple linear regression example illustrating the point that adding covariates that are independent of the response variable can increase the ability of the model to detect a correlated variable. This is not only a problem of power to detect the relationship, but can actually decrease the model's ability to assign lower p-values to variables that are actually correlated with the response variable. Code to reproduce the example is provided here, as well as a summary of the results.

```
for (k in c(1, 10, 25, 50, 75, 95)) {
    sp1 <- rnorm(n = samp, 0, 1)
    sp2 <- sp1 + rnorm(n = samp, 0, 2)
    sp3 <- rnorm(samp, 0, 1)

    cov <- matrix(data = rnorm(samp * k, 0, 1), nrow = samp, ncol = k)

    data <- data.frame(sp1 = sp1, sp2 = sp2, sp3 = sp3)
    data <- cbind(data, cov)

    model <- lm(sp1 ~ ., data)
    model_summary <- data.frame(summary(model)$coefficients)

    if (model_summary[["Pr...t.."]][3] < model_summary[["Pr...t.."]][2] {
        print("mistake")
    }
```

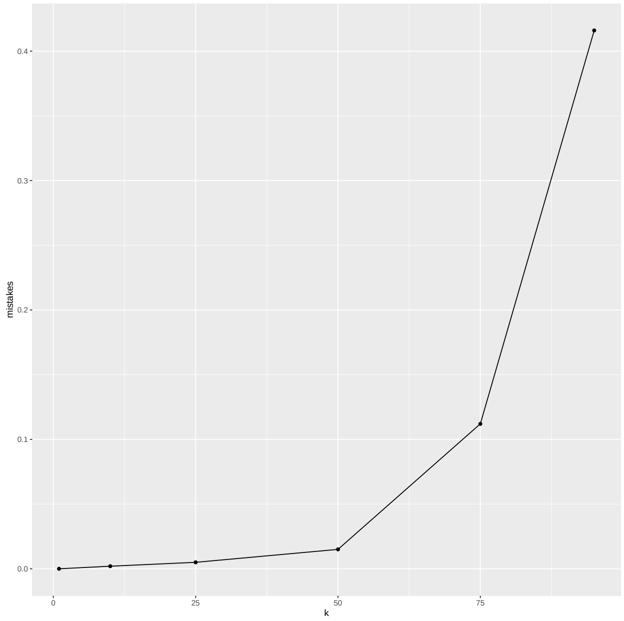


Figure S17: Even when all linear regression modeling assumptions are met, adding covariates can decrease modeling accuracy. k is the number of covariates and mistakes is the rate at which the linear regression p-value for a variable uncorrelated to the response variable is less than the p-value for a correlated variable.

}

### I Multicollinearity

According to ROC curves, we find that including environmental covariates decreases the model performance for set-parameter ecological simulations data but doesn't affect the model performance for Random-parameter ecological simulations data. Multicollinearity is unrelated to Type I error given the appropriate FDR control, but increases Type II error [5]. Thus, multicollinearity between species and covariates in set-parameter ecological simulations data is high and is at least partly responsible for the decrease in model performance when covariates are included. The multicollinearity of random-parameter ecological simulations data is low which doesn't affect the model performance when covariates are included.

Here, we show the variance inflation factor (VIF), a metric of collinearity, for 100 simulations of 10 species with 10,000 samples each for both ecological simulation sets. For set-parameter simulation the mean of the VIF values is 12.26 while for Random-parameter the mean of the VIF values is 2.44. VIF is greater than 9 is generally considered indicative of high multicollinearity. We therefor assume that multicollinearity is, at least in part, the cause of the observed decrease of model performance presented in the ROC curves.

We note that adding more covariates can decrease the performance of models even if variables are all statistically independent H. However, we can see this as multicollinearity that arises from correlation in samples in spite of variables being independent.

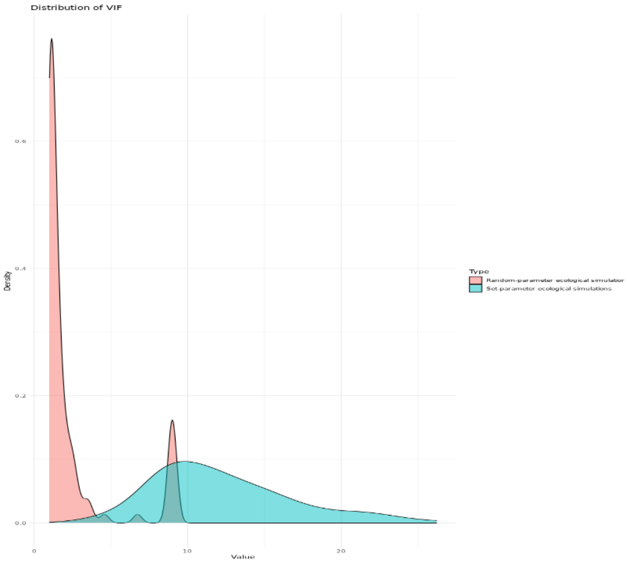


Figure S18: VIF values for 100 simulations using ecological simulations using random parameters and 100 simulations using a single parameter set, both with 10 species and 10 covariates.

#### J SPIEC-EASI performance on low number of species

When low numbers of species were used, SPIEC-EASI is unable to infer interactions with positive correlation. In fact, it seems to preferentially infer interactions between non-interacting species over those that interact (correlate) positively. This causes a strange shape of the ROC curve when all interactions are included. This does not occur for larger numbers of species (meeting the assumption of the method). This observation holds for both ecological and covariance matrix simulations.

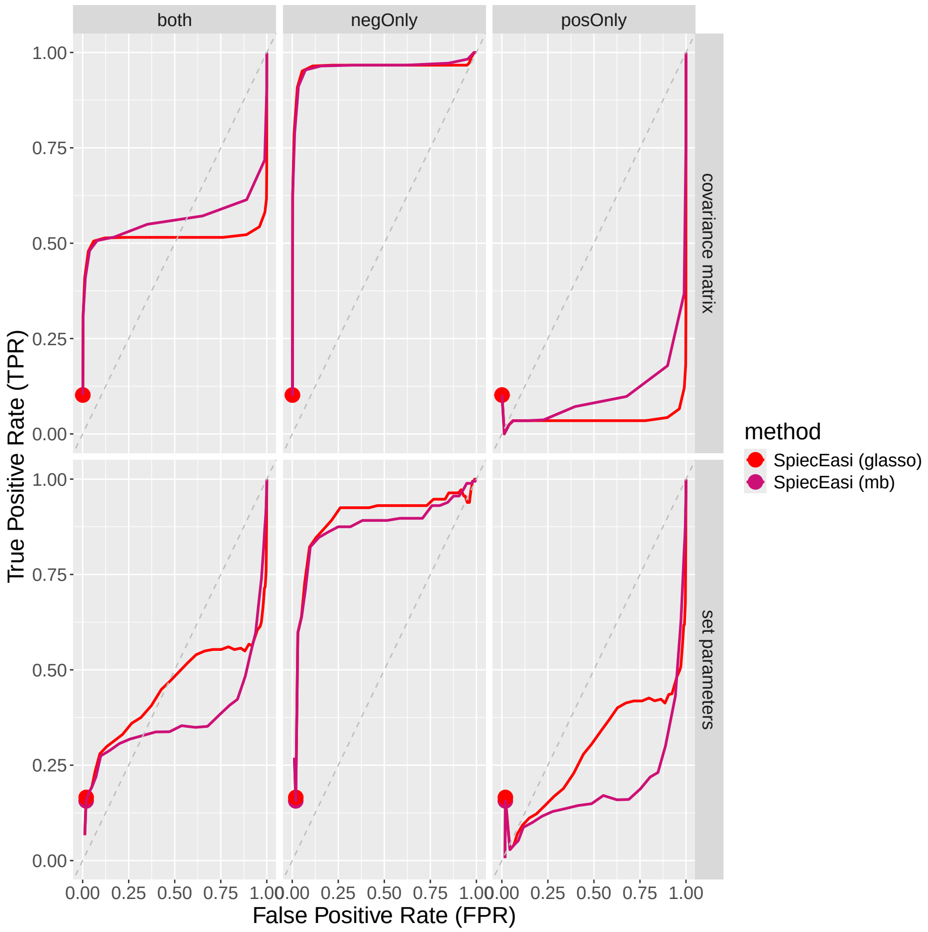


Figure S19: ROC plots for SPIEC-EASI for simulated data with 10 species when the actual interactions were limited to positive interactions only or negative interactions only, in comparison to the default where both directions are included.

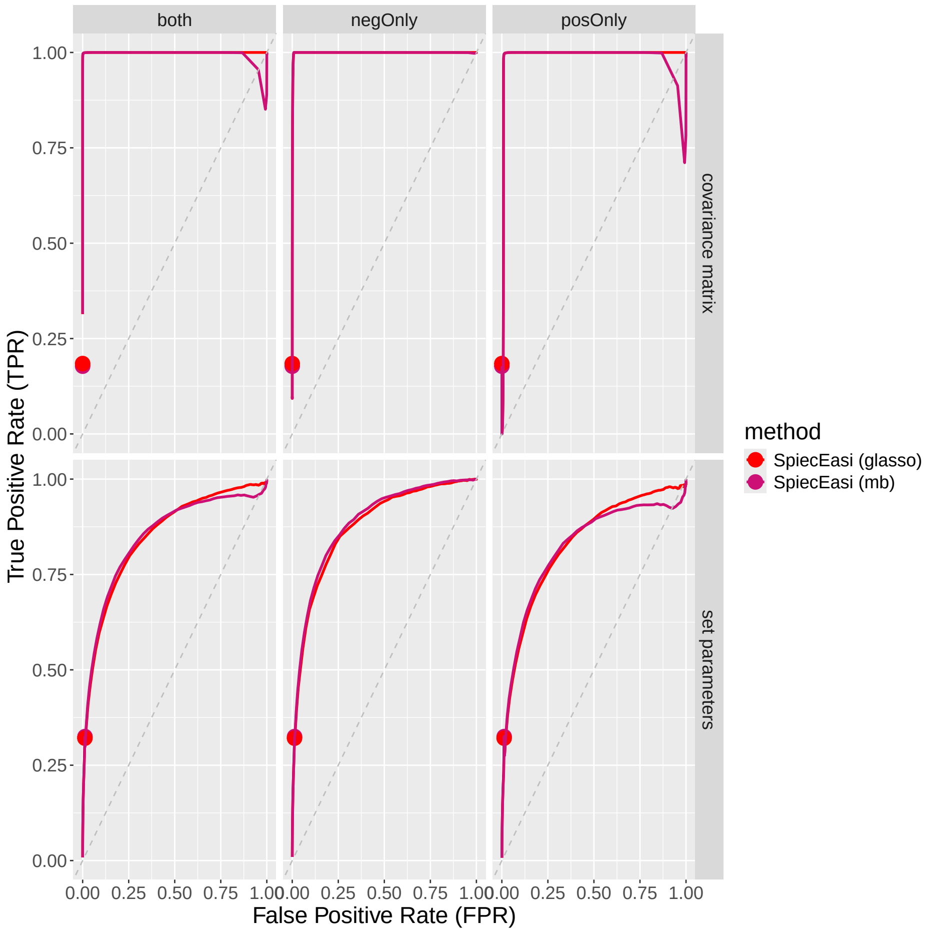


Figure S20: ROC plots for SPIEC-EASI for simulated data with 100 species when the actual interactions were limited to positive interactions only or negative interactions only, in comparison to the default where both directions are included.

### K Random forest performance on test set

| | Logistic 10,000 samples | Logistic 100 samples | Linear 10,000 samples | Linear 100 samples |
|---|---|---|---|---|
| Naive RMSE | 0.1588 | 0.3165 | 0.1820 | 0.3872 |
| Random Forest RMSE | 0.1505 | 0.2844 | 0.1492 | 0.3510 |
| Improvement | 0.052 | 0.101 | 0.180 | 0.094 |

Table 1: Predictive success of random forest for FDR $\sim$ simulation parameters. FDR was for for direct, symmetric interactions, analysis performed with 10 species in *ecological simulations* with random parameter settings and 10,000 samples per simulation. Used Benjamini–Hochberg correction with FDR control level of 0.05 for linear and logistic regression of each species as a function of all other species data. Naive estimator is the median of the observed false discovery rates. All analyses performed without covariates. Random forest trained on 1000 simulations. Test set is 100 simulations.

| | Logistic 10,000 samples | Logistic 100 samples | Linear 10,000 samples | Linear 100 samples |
|---|---|---|---|---|
| Naive RMSE | 0.2318 | 0.3909 | 0.2693 | 0.5 |
| Random Forest RMSE | 0.2135 | 0.3385 | 0.2345 | 0.4229 |
| Improvement | 0.079 | 0.134 | 0.130 | 0.154 |

Table 2: Predictive success of random forest for FDR $\sim$ simulation parameters. FDR was for for direct, symmetric interactions, analysis performed with 10 species in *ecological simulations* with random parameter settings and 10,000 samples per simulation. Used Benjamini–Hochberg correction with FDR control level of 0.05 for linear and logistic regression of each species data as a function of all other species data. Naive estimator is the median of the observed false discovery rates. All analyses performed without covariates. Random forest trained on 1000 simulations. Test set is 100 simulations.

### L Random forest ICE plots

ICE plots show the influence each predictor has on the response variable in a trained random forest.

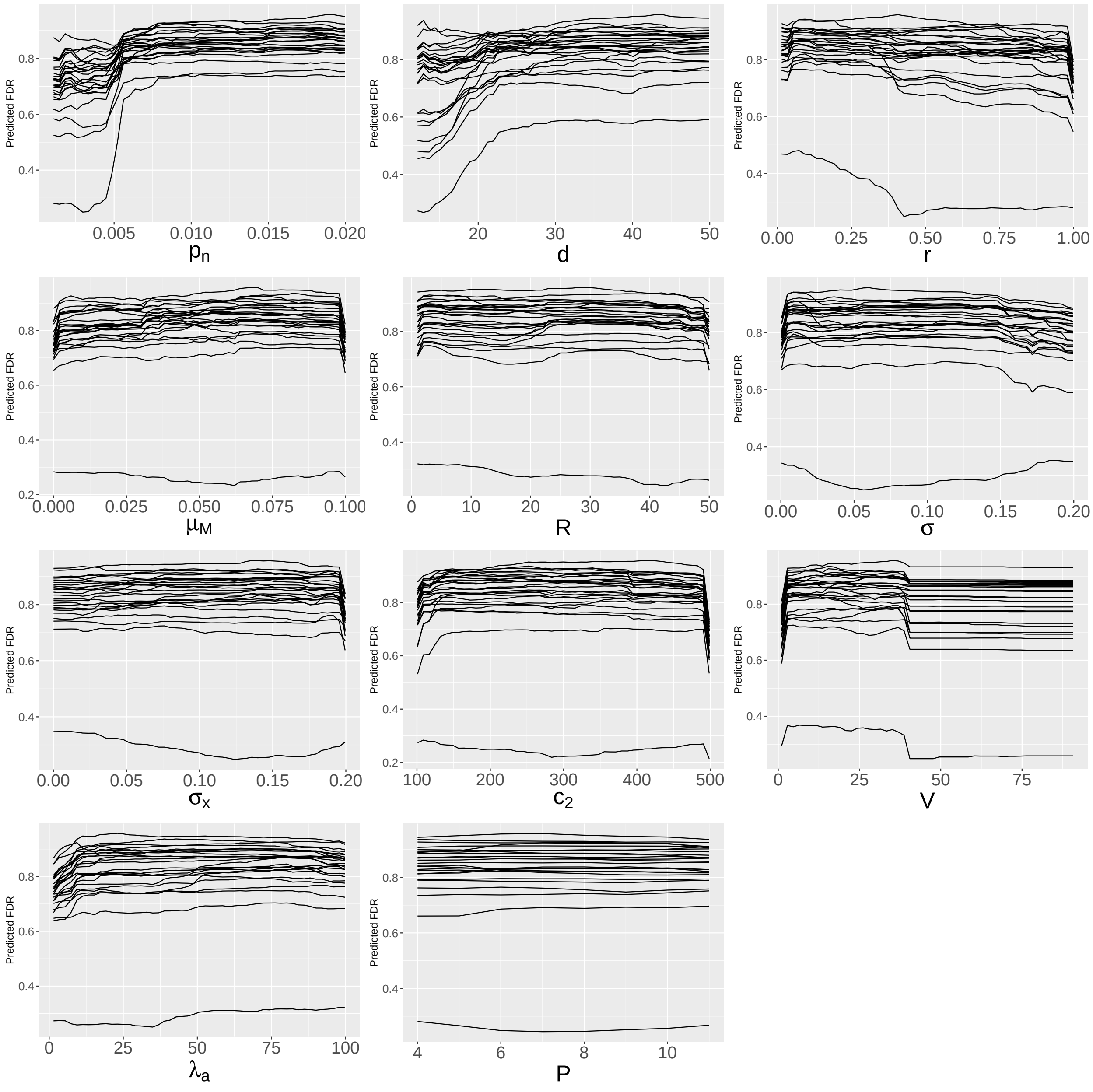


Figure S21: ICE plots for random forest predictors for FDR $\sim$ simulation parameters. FDR was for for direct, symmetric interactions; analysis performed with 10 species in *ecological simulations* with random parameter settings. Used Benjamini–Hochberg correction with FDR control level of 0.05 for linear and logistic regression of each species data as a function of all other species data. Analysis performed with logistic regression with 10,000 samples per simulation and random forest was trained on results from 1,000 simulations with 10 species.

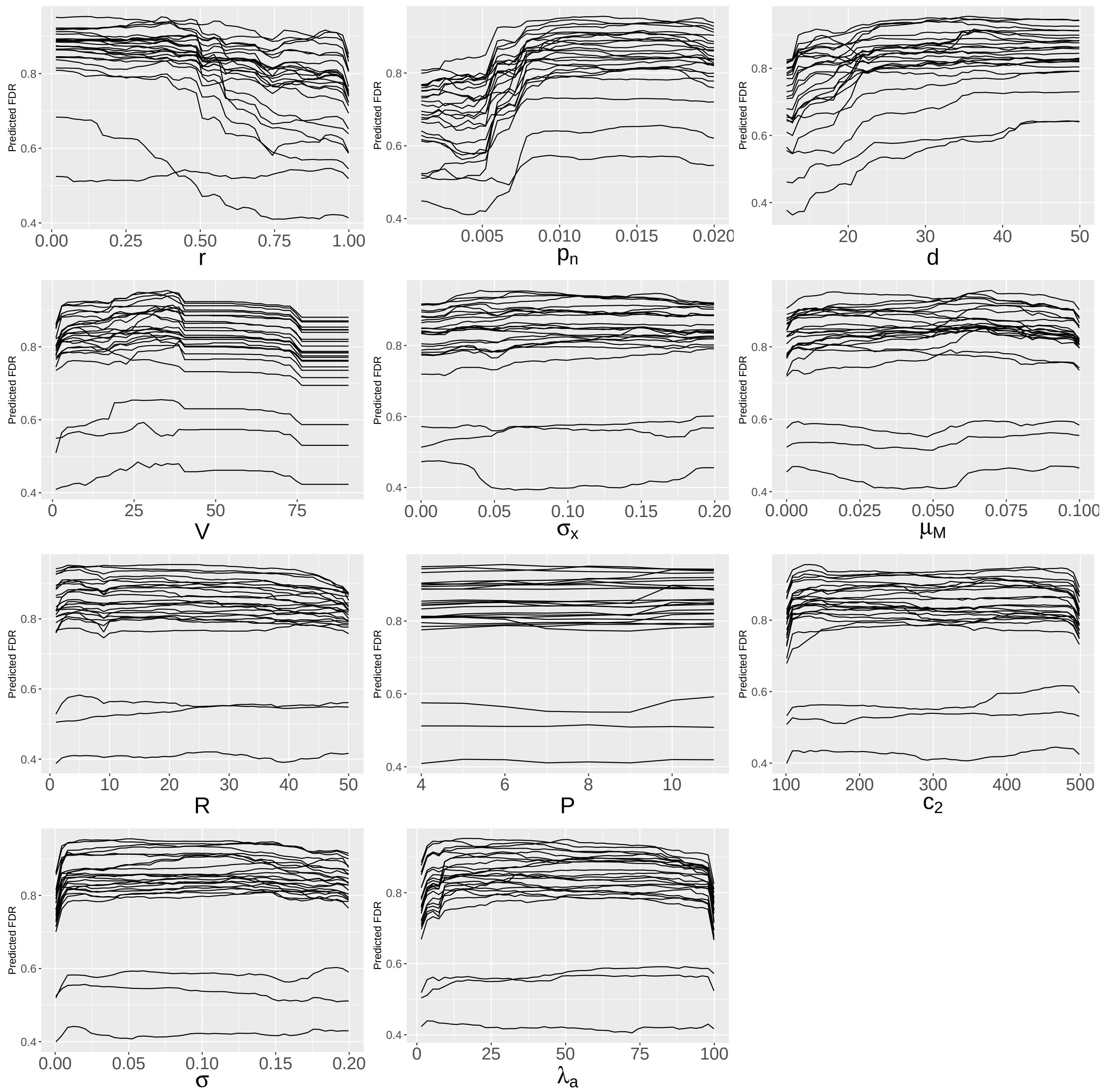


Figure S22: ICE plots for random forest predictors for FDR ∼ simulation parameters. FDR was for for direct, symmetric interactions; analysis performed with 10 species in *ecological simulations* with random parameter settings. Used Benjamini–Hochberg correction with FDR control level of 0.05 for linear and logistic regression of each species data as a function of all other species data. Analysis performed with linear regression with 10,000 samples per simulation and random forest was trained on results from 1,000 simulations with 10 species.

### M Actual interactions cause higher percent presence on average in my simulation, but logistic regression infers interactions more for species with close to 50% presence

We observed that in some of the simulations, in particular those with higher numbers of species, the species that interacted with other species tended to have very low or very high abundance. This is an emergent property of the simulation model that may or may not be reproduced in real data.

We also observe that, independently of whether there were actual interactions, logistic regression is more likely to infer interactions between species with intermediate abundance. We hypothesize that this is because the power to infer interactions is highest when the percent presence is around 50%. This is intuitive if you consider a species that is never present or a species that is always present in the data. There will be no way to see an interaction with that species using detection/non-detection data. This effect will not be as pronounced in abundance data but it may still have an effect if there are a lot of 0's for some species, or when the read abundances have low variance.

Although we did not test this effect for other methods, we believe it may have some influence since the pattern in the data will hold regardless.

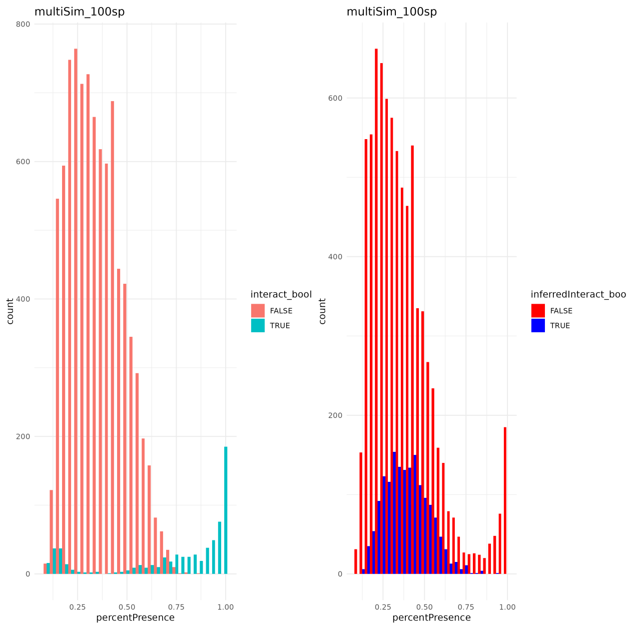


Figure S23: Distribution of percent presence of species in the set-parameter data, split by whether an interaction exists (left) and whether an interaction is inferred (right). For 100 species, set parameters, logistic regression noCov, p-value cutoff 0.9999, 100 samples

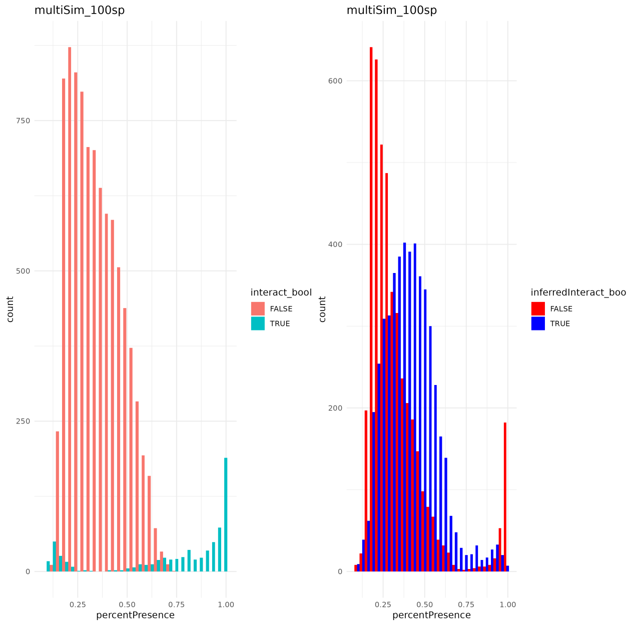


Figure S24: Distribution of percent presence of species in the set-parameter data, split by whether an interaction exists (left) and whether an interaction is inferred (right). For 100 species, set parameters, logistic regression noCov, p-value cutoff $1 * 10^{-16}$, 10,000 samples

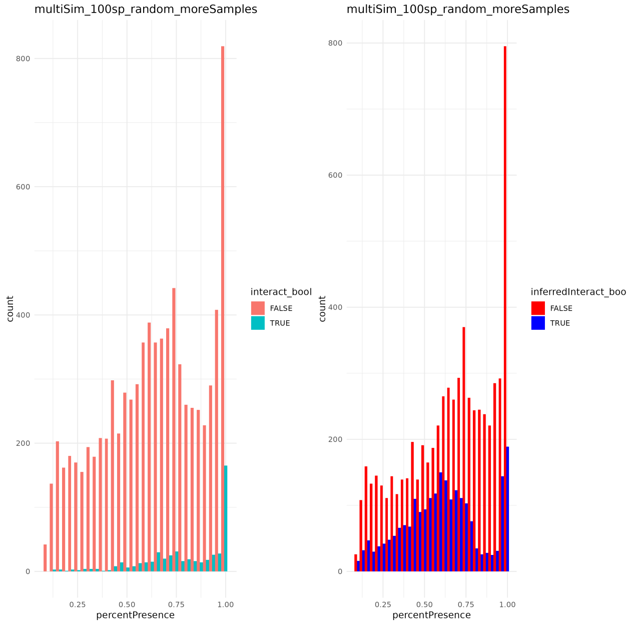


Figure S25: Distribution of percent presence of species in the set-parameter data, split by whether an interaction exists (left) and whether an interaction is inferred (right). For 100 species, random parameters, logistic regression noCov, p-value cutoff 0.9999, 100 samples

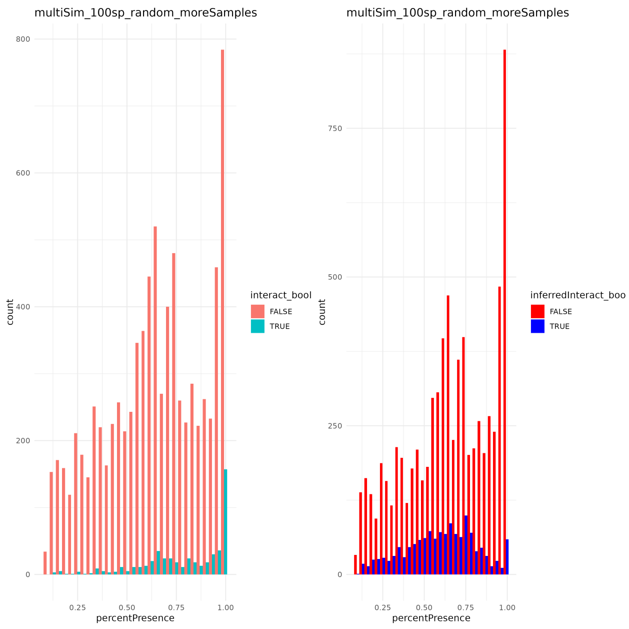


Figure S26: Distribution of percent presence of species in the set-parameter data, split by whether an interaction exists (left) and whether an interaction is inferred (right). For 100 species, random parameters, logistic regression noCov, p-value cutoff $1 * 10^{-8}$, 10000 samples

#### N Distribution of covariances for *Covariance Matrix Simulations*

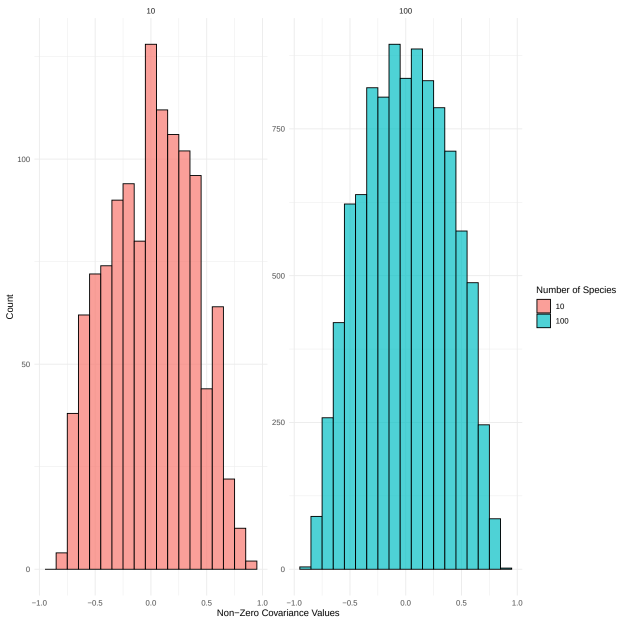


Figure S27: Distribution of target covariances in the covariance matrix simulation model without covariates.

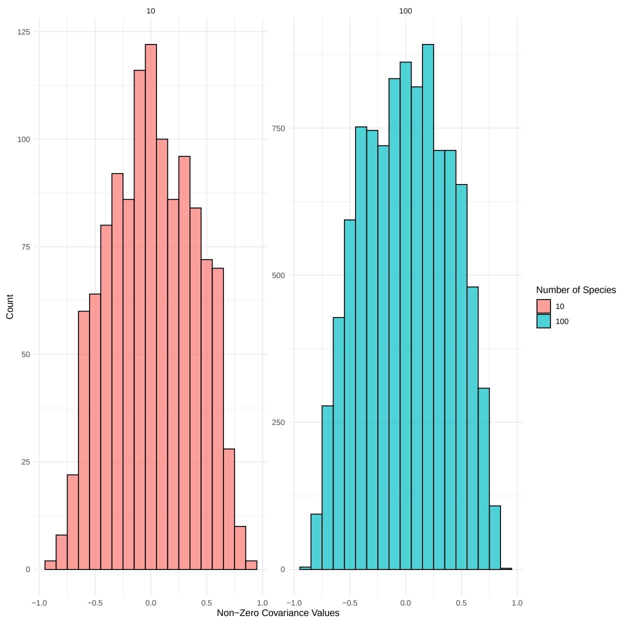


Figure S28: Distribution of target covariances in the covariance matrix simulation model with covariates.

### O Comparison of BH and Bonferroni correction

We also tested how the results changed by using a Bonferroni correction for the linear and logistic regression methods (instead of Benjamini-Hochberg correction, or uncorrected 0.05 cutoff). The overall trends in the results were similar. Uncorrected is the least conservative, Benjamini-Hochberg correction [6] is intermediate, and Bonferroni is the most conservative. All results shown are for direct, symmetric interactions.

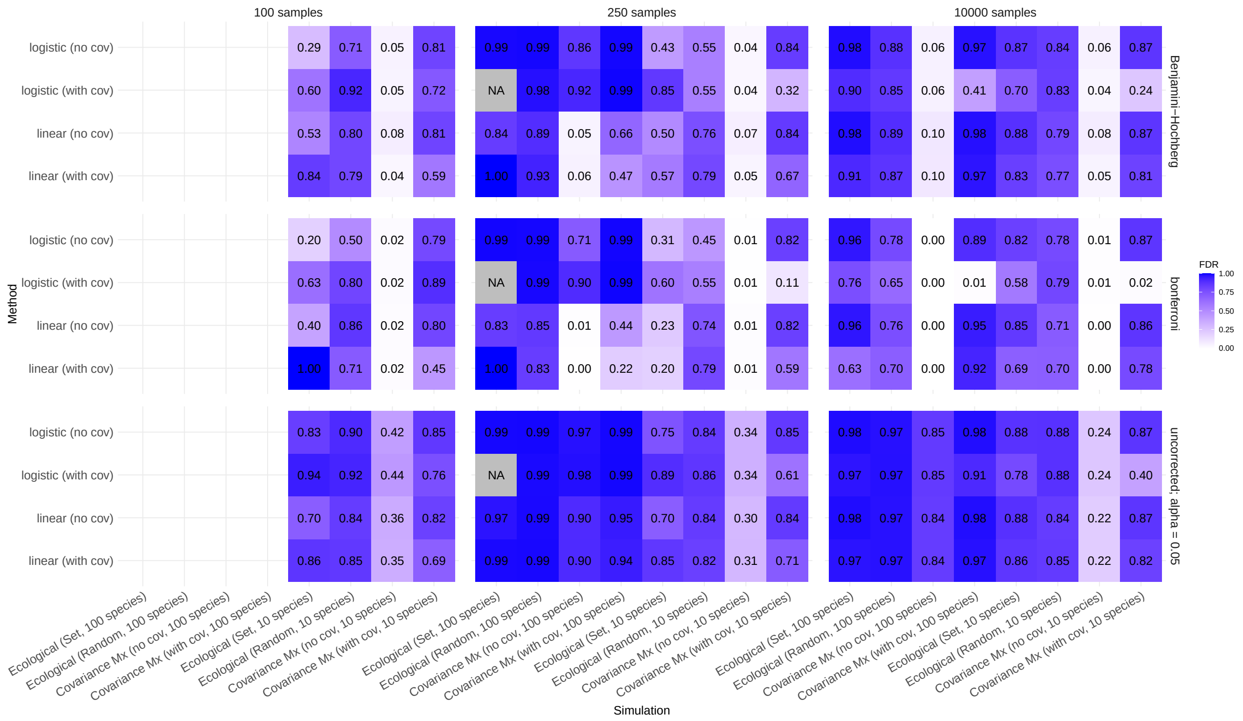


Figure S29: Distribution of target covariances in the covariance matrix simulation model with covariates.

### P Linear regression on Gaussian z-values

For the Poisson reads in the *covariance matrix simulation*, we see that no method performs well when covariates are included. We concluded from this that none of the methods can successfully correct for covariates with the violations of modeling assumptions presented by the Poisson distributed reads. In further support of this conclusion, we ran linear regression on the latent Gaussian z-values in the model rather than the Poisson distributed counts derived from these values (see model description of *covariance matrix simulation* in the main text; $z_j(s,t)$). We see that linear regression performs well when its assumptions are met better, even with covariates included in the simulation, as long as the covariates are also accounted for in the regression. When covariates are included in the simualtion but not corrected for in the regression, as expected we still see poor performance due to confounding variables.

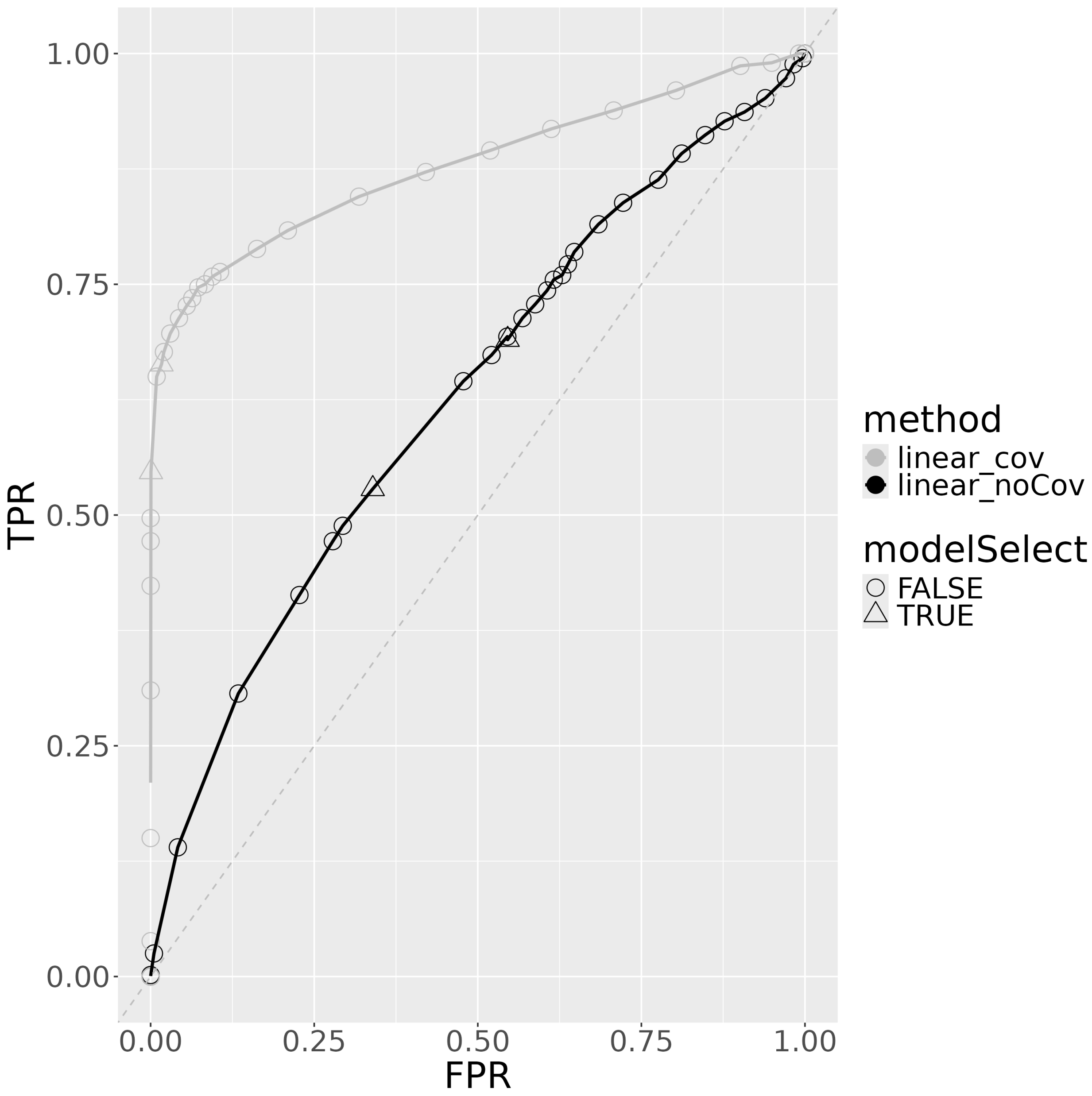


Figure S30: ROC curve for linear regression performed on 250 samples of the intermediate Gaussian latent variable $z$ from the covariance matrix simulations with covariate effects included in the simulation. Linear_cov refers to the linear regression with covariates included as regressors, whereas linear_noCov refers to linear regression with only the other species included as regressors. Triangles represent FDR corrected values (Bonferroni or BH).

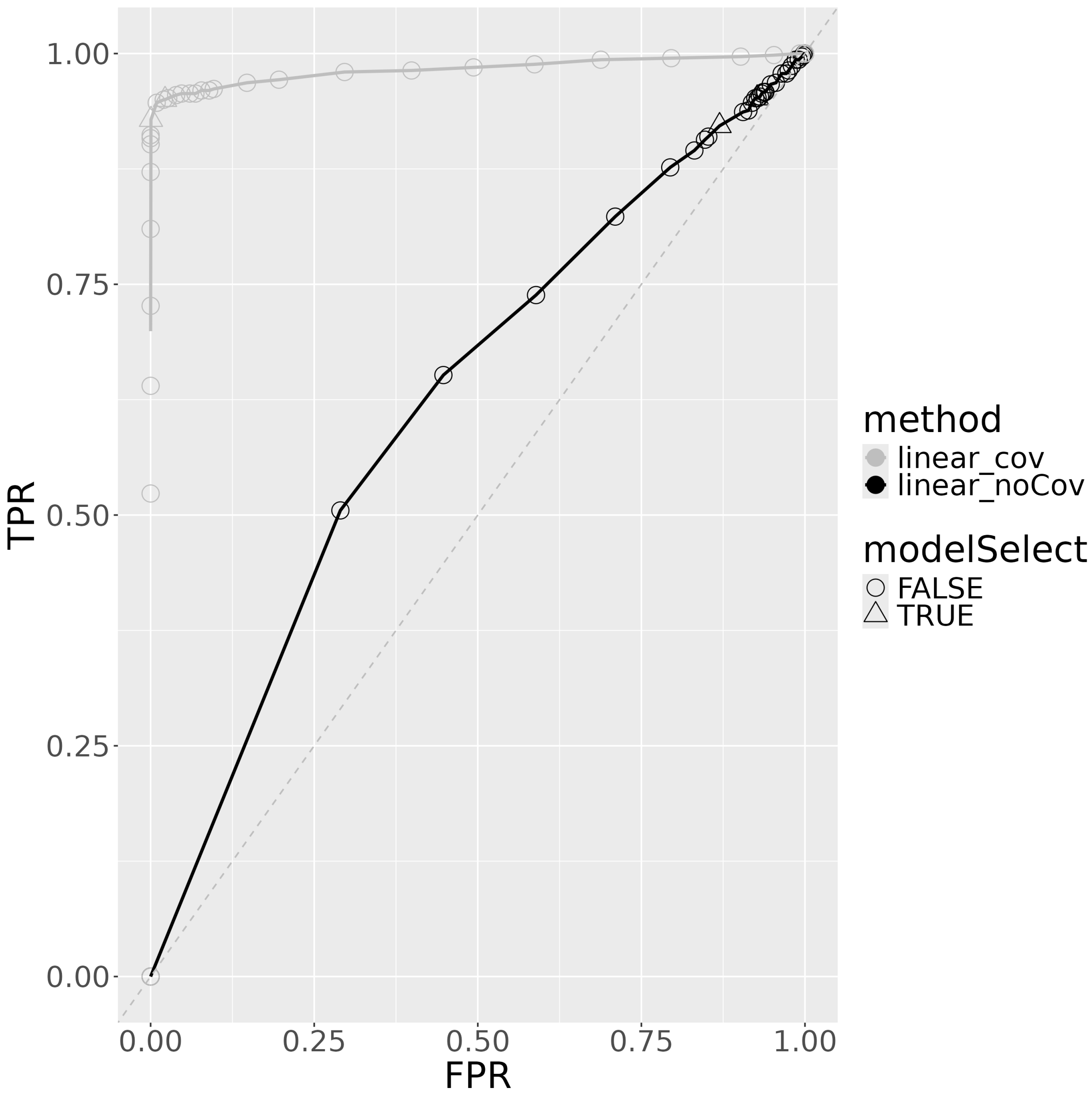


Figure S31: ROC curve for linear regression performed on 10000 samples of the intermediate Gaussian latent variable $z$ from the covariance matrix simulations with covariate effects included in the simulation. Linear_cov refers to the linear regression with covariates included as regressors, whereas linear_noCov refers to linear regression with only the other species included as regressors. Triangles represent FDR corrected values (Bonferroni or BH).
